# The Genomic Architecture of a Rapid Island Radiation: Recombination Rate Variation, Chromosome Structure, and Genome Assembly of the Hawaiian Cricket *Laupala*

**DOI:** 10.1101/160952

**Authors:** Thomas Blankers, Kevin P. Oh, Aureliano Bombarely, Kerry L. Shaw

## Abstract

Phenotypic evolution and speciation depend on recombination in many ways. Within populations, recombination can promote adaptation by bringing together favorable mutations and decoupling beneficial and deleterious alleles. As populations diverge, cross-over can give rise to maladapted recombinants and impede or reverse diversification. Suppressed recombination due to genomic rearrangements, modifier alleles, and intrinsic chromosomal properties may offer a shield against maladaptive gene flow eroding co-adapted gene complexes. Both theoretical and empirical results support this relationship. However, little is known about this relationship in the context of behavioral isolation, where co-evolving signals and preferences are the major hybridization barrier. Here we examine the genomic architecture of recently diverged, sexually isolated Hawaiian swordtail crickets (*Laupala*). We assemble a *de novo* genome and generate three dense linkage maps from interspecies crosses. In line with expectations based on the species’ recent divergence and successful interbreeding in the lab, the linkage maps are highly collinear and show no evidence for large-scale chromosomal rearrangements. The maps were then used to anchor the assembly to pseudomolecules and estimate recombination rates across the genome. We tested the hypothesis that loci involved in behavioral isolation (song and preference divergence) are in regions of low interspecific recombination. Contrary to our expectations, a genomic region where a male song QTL co-localizes with a female preference QTL was not associated with particularly low recombination rates. This study provides important novel genomic resources for an emerging evolutionary genetics model system and suggests that trait-preference co-evolution is not necessarily facilitated by locally suppressed recombination.

## INTRODUCTION

Speciation is contingent on the accumulation of genomic variation and the formation of barriers that prevent gene flow between populations. Genomes diverge under the influence of selection and drift, while gene flow counteracts this divergence by homogenizing the genome (Felsenstein 1981; Kirkpatrick and Ravigne 2002; Gavrilets 2003). To appreciate the speciation process and the origin of the fascinating diversity of life on earth, we need to understand the interaction between the mechanisms that change allele frequencies and the mechanisms that govern the association of beneficial and deleterious alleles with other alleles. A key process in this interaction is recombination, which creates new allelic combinations during meiosis in sexually reproducing organisms.

Any association between loci that underlie environmental adaptation or between loci underlying co-evolving (sexual) signals and signal responses (i.e. co-adapted gene complexes) will be affected by recombination (Felsenstein 1981). Within populations, recombination can mitigate Hill-Robertson interference by combining locally adaptive alleles from different genomic backgrounds and by decoupling beneficial and deleterious alleles (Hill and Robertson 1966); recombination can also influence the covariance between sexual traits and preference across sexes (Smith and Haigh 1974; Smith 1978; Gillespie 2000; Otto 2009). As such, recombination might increase the efficiency of background selection (purging deleterious alleles), sexual selection (through signal-preference co-evolution), and local adaptation in the earliest stages of speciation (by linking locally adapted alleles; Noor *et al.* 2001; Rieseberg 2001; Kirkpatrick and Ravigne 2002; Yeaman and Whitlock 2011).

Between divergent populations with some (but incomplete) reproductive isolation, recombination can also counteract population divergence and prevent the closure of a reproductive boundary by creating combinations of alleles that are favorable in different contexts (Noor *et al.* 2001; Rieseberg 2001; Coyne and Orr 2004; Ortiz-Barrientos *et al.* 2016). It is important to realize that interspecific recombination is constrained both by intrinsic properties of the species’ genomes that also constrain intraspecific recombination, as well as by the effects from (divergent) selection and alternatively fixed chromosomal rearrangements (Yeaman and Whitlock 2011; Feder *et al.* 2012). The most intensely studied chromosomal rearrangements suppressing recombination between divergent populations are inversions. Inversions can suppress recombination locally in the genome and, thus, promote reproductive isolation, by trapping genetic incompatibilities in linkage blocks (Noor *et al.* 2001), acting synergistically with other genes causing isolation (Rieseberg 2001), or by linking locally adaptive alleles (Kirkpatrick and Barton 2006). Other chromosomal rearrangements, such as translocations and transposable elements, can likewise contribute to ‘chromosomal speciation’ (Rieseberg 2001) as well as to preventing gene flow and furthering genetic divergence among heterospecifics.

Interestingly, there is ubiquitous among-species variation in recombination rates (Wilfert *et al.* 2007; Smukowski and Noor 2011). In insects, for example, rates vary from 16.1 cM/Mb (centi-Morgans per megabase) in *Apis melifera* to 0.1 cM/Mb in the mosquito *Armigeres subalbatus* (Wilfert *et al.* 2007). There is also variation across the genome within individuals. For example, 50-fold differences have been observed within single chromosomes of humans and birds (Myers *et al.* 2005; Singhal *et al.* 2015). These patterns of variation underline that the efficacy of selection acting within species may differ across taxa and across genomes of the same species.

A major prediction following from theoretical work is that favorable allele combinations that promote ecological adaptation are more likely to reside in regions of low recombination. Recombination frustrates natural selection by breaking up associations between segregating alleles that are locally adaptive within the resident population and counteracts divergent selection if there is gene flow between recently diverged populations (Bürger and Akerman 2011; Yeaman and Whitlock 2011; Yeaman 2013). So far, empirical evidence for the prediction that locally adaptive alleles reside in regions of low recombination is not conclusive (Roesti *et al.* 2013; Burri *et al.* 2015; Marques *et al.* 2016). However, a recent study indicated that the interaction between gene flow and divergent selection is a strong predictor for the association between adaptive alleles and regions of low recombination in multiple species of stickleback fish (Samuk *et al.* 2017).

However, it is unclear how these predictions apply to the evolution of behavioral isolation. Theoretical models of speciation by sexual selection depend on linkage disequilibrium between sexual signaling traits and corresponding preference genes (Fisher 1930; Lande 1981; Kirkpatrick 1982). Linkage disequilibrium between trait and preference genes can come about by assortative mating (Lande 1981; Andersson and Simmons 2006) or by physical linkage (Kirkpatrick and Hall 2004), either through closely linked loci or through pleiotropy (a single gene affecting both signal and preference phenotypes). Here, the role of recombination is more complex: On the one hand, recombination can help consolidate loci brought together by nonrandom mating and as such facilitate linkage disequilibrium between trait and preference (Kirkpatrick and Ravigne 2002). On the other hand, recombination can also tear apart co-adapted trait and preference alleles if genes are exchanged between populations that differ in mating phenotypes. Therefore, recombination between sexually divergent populations in sympatry and parapatry often compromises differentiation in mating phenotypes and hinders speciation (Arnegard *et al.* 2004; Servedio 2009, 2015; Servedio and Burger 2014). However, there has been limited empirical insight into the relationship between trait-preference co-evolution and genome-wide variation in recombination rates (see Davey *et al.* 2017 for a recent exception).

Here, we examine the genomic architecture, specifically structural variation and heterogeneity in interspecific recombination, of four closely related, sexually isolated species of Hawaiian swordtail crickets from the genus *Laupala*. *Laupala* is one of the fastest speciating taxa known to date (Mendelson and Shaw 2005). The 38 morphologically cryptic species, each endemic to a single island of the Hawaiian archipelago (Otte 1994; Shaw 2000a) are the product of a recent evolutionary radiation. Evidence suggests that speciation by sexual selection on the acoustic communication system has driven this rapid diversification, as both male mating song and female acoustic preferences have diverged extensively among *Laupala* species (Otte 1994; Shaw 2000b; Mendelson and Shaw 2002). Sexual trait evolution strongly contributes to the onset and maintenance of reproductive isolation (Mendelson and Shaw 2002; Grace and Shaw 2011). Quantitative variation in one key temporal property of male song (pulse rate) and corresponding female preference strongly covaries across species and across populations within species (Shaw 2000b; Grace and Shaw 2011). Although the mechanisms of trait-preference co-evolution require further study, there is evidence that both are associated with a polygenic basis and that genetic loci controlling quantitative variation in traits and preferences are physically linked in the genome (Shaw and Lesnick 2009; Wiley *et al.* 2012). Notably, one of the major song quantitative trait loci (QTL; haploid effect size ∼ 9%) co-localizes with the first mapped preference QTL (haploid effect size ∼ 14%). Directional effects of song QTL provide additional evidence that (sexual) selection is driving divergence between species (Shaw *et al.* 2007).

The species pairs involved in this study, *L. kohalensis* and *L. pruna*, and *L. paranigra* and *L. kona*, are endemic to the Big Island, the youngest island of the Hawaiian archipelago (Fig 1A, B). Although these species pairs have apparently diverged in allopatry within the Big Island, past or future migration is likely, given their geographical proximity. Indeed, although allopatric and more closely related to *L. kohalensis, L. pruna* currently overlaps in distribution with *L. paranigra* (Fig 1B). The discordance between nuclear and mitochondrial phylogenies (Shaw 2002) and the limited degree of postzygotic isolation between some species pairs further emphasize the possibility of gene flow across natural populations. Together, the biogeography and the genetics of song and preference variation in this system provide a unique opportunity to explore the interaction between interspecific recombination rate variation, co-evolution of mating traits, and speciation.

**Figure 1.**
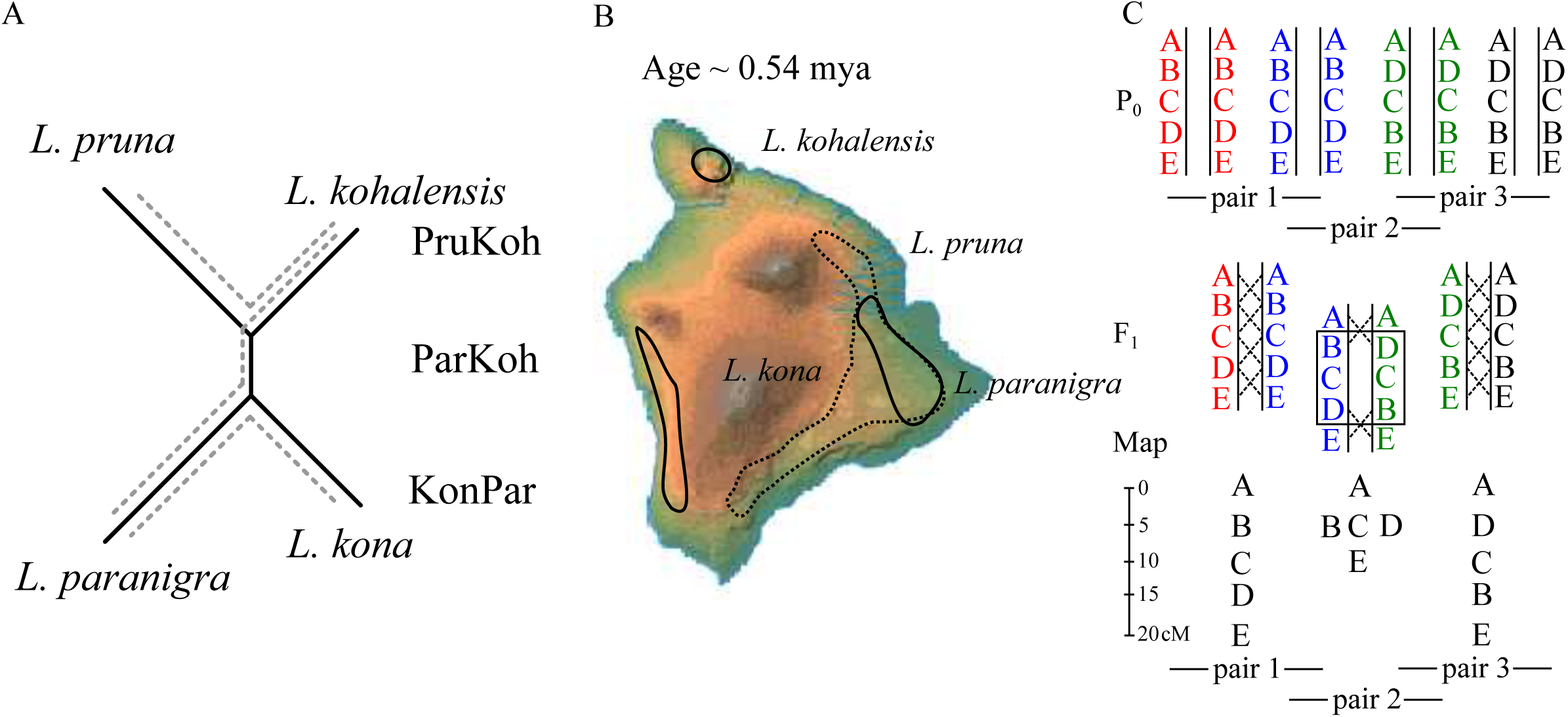
Study design. (A) The phylogenetic relationships of studied *Laupala* species based on a neighbor joining tree generated from genetic distances among the parental lines used in this study. Dashed grey lines connect species pairs that were crossed. (B) Approximate distributions of the studied species on the Big Island of Hawaii. (C) Hypothetical segregation and linkage map construction for five genetic loci A, B, C, D, and E in three crosses of fours species. The genetic distance between the loci is 5 centi-Morgan (cM) in each of the four species. Loci [B,C,D] are inverted in the green and black species. When two species that have alternative karyotypes for the inversion are crossed (pair 2), loci in the inversion will not recombine in the first generation hybrid, resulting in reduced genetic (map) length in the second generation hybrid. Other chromosomal rearrangements will have similar effects. Only if two crosses involve homokaryotypic species pairs that have alternative karyotypes can an inversion be detected in a comparison of intercross linkage maps.

We first assemble a *de novo L. kohalensis* draft genome and then obtain thousands of SNP markers for heterogeneously hybrid offspring from three laboratory-generated interspecific crosses. We then generate three dense linkage maps and compare these maps to test the hypothesis that the genomic architectures of young, sexually differentiated species are largely collinear (similar marker order) and have conserved interspecific recombination frequencies (similar marker distances). There is some variation in the level of overall differentiation in the species pairs studied here, but all lineages are young (approximately 0.5 million years or less, Fig 1). It is commonly expected that strong prezygotic isolation can evolve rapidly and largely in the absence of intrinsic postzygotic isolating mechanisms (Coyne and Orr 2004), but explicit comparisons of chromosomal architectures across behaviorally isolated species are rare. We compare the maps visually and use variation in maker order and length (measured in genetic distance, or centi-Morgans [cM]) as indicators of possible chromosomal rearrangements affecting the recombination rates differently in different crosses (Fig 1C). Then, from the large amount of information on linkage across many genomic markers from three hybrid crosses, we anchor the draft genome assembly to pseudomolecules and estimate the landscape of recombination across the genome. Finally, using an additional map that integrates the amplified fragment length polymorphism (AFLP) markers from previous QTL studies in *L. kohalensis* and *L. paranigra*, we approximate the location of known male song QTL, including one co-localizing with a female acoustic preference QTL, on the pseudomolecules. We examine local variation in recombination rates across the genome and in relation to the location of the song and preference QTL to test the hypothesis that song-preference co-evolution is facilitated by suppressed interspecific recombination. This study provides important insight into the role of the genomic architecture during divergence of closely related species separated by premating barriers.

## MATERIAL & METHODS

### De novo genome assembly

The *Laupala kohalensis* draft genome (estimated genome size ∼ 1.9 Gb; Petrov *et al.* 2000) was sequenced using the Illumina HiSeq 2500 platform. DNA was isolated with the DNeasy Blood & Tissue Kits (Qiagen Inc., Valencia, CA, USA) from six immature female crickets (c. five months of age) chosen randomly from a laboratory stock population (approximate lab generation=14). Females were chosen to balance DNA content of sex chromosomes to autosomes (female crickets are XX; male crickets are XO). DNA was subsequently pooled for sequencing. Four different libraries were created: a paired-end library with an estimated insert size of 200 bp (sequenced by Cornell Biotechnology Resource Center), a paired-end library with an estimated insert size of 500 bp, and two mate-pair libraries with insert sizes of 2 and 5 Kb (sequenced by Cornell Weill College Genomics Resources Core Facility).

Reads were processed using Fastq-mcf from the Ea-Utils package (Aronesty 2011) with the parameters -q 30 (trim nucleotides from the extremes of the read with qscore below 30) and -l 50 (discard reads with lengths below 50 bp). Read duplications were removed using PrinSeq (Schmieder and Edwards 2011) and reads were corrected using Musket with the default parameters (Liu *et al.* 2013).

Reads were assembled using SoapDeNovo2 (Luo *et al.* 2012). The reads were assembled using different Kmer sizes (k = 31, 39, 47, 55, 63, 71, 79 and 87). The 87-mer assembly produced the best assembly (based on N50/L50, assembly size, and number of scaffolds). Scaffolds and contigs were renamed using an in-house Perl script. Gaps were filled using GapCloser from the SoapDeNovo2 package.

The gene space covered by the assembly was evaluated using three different approaches. (1) *Laupala kohalensis* unigenes produced by the Gene Index initiative (Cricket release 2.0: http://compbio.dfci.harvard.edu/cgi-bin/tgi/gimain.pl?gudb=cricket) were mapped using Blat (Kent 2002). Only unigenes mapping with 90% or more of their length were considered; (2) 50 bp paired-end RNA-seq reads from a congeneric species, *L. cerasina* were mapped using Tophat2 (Kim *et al.* 2013). Reads were processed using the same methodology described above, but using a minimum length of 30 bp; (3) using BUSCO (Simão *et al.* 2015) to search for conserved eukaryotic and arthropod genes.

### Samples

We generated three F_2_ interspecies hybrid families to estimate genetic maps. Multiple F_1_ male and sibling females were intercrossed to generate F_2_ mapping populations for the following species crosses: (1) a *L. kohalensis* female and *L. paranigra* male (“ParKoh", 178 genotyped F_2_ hybrid offspring; previously reported in Shaw et al. 2007); (2) a *L. kohlanesis* female and a *L. pruna* male (“PruKoh", 193 genotyped F_2_ hybrid offspring); (3) a *L. paranigra* female and a *L. kona* male (“KonPar", 263 genotyped F_2_ hybrid offspring). These four species are part of a recently radiated clade showing conspicuous mating song divergence (Mendelson and Shaw 2005). Approximate geographic distributions of the species, phylogenetic relationships and parent collection localities are shown in Fig 1 and in Table S1. Crickets used in crosses were a combination of lab stock and outbred individuals (*L. kohalensis* [for ParKoh] and *L. paranigra* [for ParKoh and KonPar] were both lab reared for 3-15 generations; *L. kohalensis* [for PruKoh], *L. pruna* and *L. kona* were wild-caught). All parental and hybrid generations were reared in a temperature-controlled room (20***°***C) on Purina cricket chow and provided water *ad libitum*.

### Genotyping

DNA was extracted from whole adults using the DNeasy Blood & Tissue Kits (Qiagen, Valencia, CA, USA). Genotype-by-Sequencing library preparation and sequencing were done in 2014 at the Genomic Diversity Facility at Cornell University following Elshire *et al.* (2011). The Pst I restriction enzyme was used for sequence digestion and DNA was sequenced on the Illumina HiSeq 2500 platform (Illumina Inc., USA).

Reads were trimmed and demultiplexed using Flexbar (Dodt *et al.* 2012) and then mapped to the *L. kohalensis de novo* draft genome using Bowtie2 (Langmead and Salzberg 2012) with default parameters. We then called SNPs using two different pipelines: The Genome Analysis Toolkit (GATK; DePristo *et al.* 2011; Van der Auwera *et al.* 2013) and FreeBayes (Garrison and Marth 2012). For GATK we used individual BAM files to generate gVCF files using ‘HaplotypeCaller’ followed by the joint genotyping step ‘GenotypeGVCF’. We then evaluated variation in SNP quality across all genotypes using custom R scripts to determine appropriate settings for hard filtering based on the following metrics (based on the recommendations for hard filtering section “Understanding and adapting the generic hard-filtering recommendations” at https://software.broadinstitute.org/gatk/ accessed on 28 February 2017): quality-by-depth, Phred-scaled *P*-value using Fisher’s Exact Test to detect strand bias, root mean square of the mapping quality of the reads, u-based z-approximation from the Mann-Whitney Rank Sum Test for mapping qualities, u-based z-approximation from the Mann-Whitney Rank Sum Test for the distance from the end of the read for reads with the alternate allele. For FreeBayes we called variants from a merged BAM file using standard filters. After variant calling we filtered the SNPs using ‘vcffilter’, a Perl library part of the VCFtools package (Danecek *et al.* 2011) based on the following metrics: quality (> 30), depth of coverage (> 10), and strand bias for the alternative and reference alleles (SAP and SRP, both > 0.0001). Finally, the variant files from the GATK pipeline and the FreeBayes pipeline were filtered to only contain biallelic SNPs with less than 10% missing genotypes using VCFtools.

We retained two final variant sets: a high-confidence set including only SNPs with identical genotype calls between the two variant discovery pipelines and the full set of SNPs which included all variants called using FreeBayes but limited to positions that were shared among the GATK and FreeBayes pipelines.

### Linkage mapping

The genotype information from the parental lines was used to assign ancestry to the SNP loci. The parents of the crosses were heterogeneously heterozygous and only ancestry informative loci were retained, i.e. all loci for which one or more of the parents was heterozygous were discarded. We were unable to obtain sequence data from the parents for PruKoh, but used sequence data from a single, non-parental *L. pruna* female, and three available *L. kohalensis* females, all from the same populations as the parents. Ancestry was inferred if all three *L. kohalensis* individuals were homozygous for one allele and the *L. pruna* individual was homozygous for the alternative allele. All other loci were discarded. The loci were then further filtered based on genotype similarity and segregation distortion (see below for details).

The linkage maps deriving from the three species crosses were generated independently and by taking a three-step approach, employing both the regression mapping and the maximum likelihood (ML) mapping functions in JoinMap 4.0 (van Ooijen 2006) as well as the three-point error-corrected ML mapping function in MapMaker 3.0 (Lander *et al.* 1987; Lincoln *et al.* 1993).

In the first step, we estimated “initial” maps that are relatively low resolution (5 cM) but with high marker order certainty. For initial maps, we first grouped (3.0 ≤ LOD ≤ 5.0) and then ordered the high-confidence markers that showed no segregation distortion (markers with χ^2^-square associated *P*-value for deviation from Mendelian inheritance < 0.05 were discarded) and for which no marker had more than 95% similarity in genotypes across individuals compared to other markers (otherwise, one of each pair was excluded). When excluding similar loci, we favored those marker loci shared among the three mapping populations over markers unique to one or two crosses. We then checked for concordance among the three mapping algorithms. In most cases, the maps were highly concordant (in ordering of the markers; with respect to cM among markers, distances differed depending on the algorithm, especially between the regression and ML methods in JoinMap). Discrepancies among the maps produced by the different algorithms for the same cross were resolved by optimizing the likelihood and total length of a given map as well as by using the information in JoinMap’s “Genotype Probabilities” and “Plausible Positions".

These initial maps were then filled out using MapMaker with marker loci passing slightly more lenient criteria: markers drawn from the full set of SNPs, with false discovery rate (Benjamini and Hochberg 1995) corrected *P* value for χ^2^-square test of deviation from Mendelian inheritance ≤ 0.05 and fewer than 99% of their genotypes in common with other markers loci. First, more informative markers (no missing genotypes, > 2.0 cM distance from other markers) were added satisfying a log-likelihood threshold of 4.0 for the positioning of the marker (i.e., assigned marker position is 10,000 times more likely than any other position in the map). Remaining markers were added at the same threshold, followed by a second round for all markers at a log-likelihood threshold of 3.0. We then used the ripple algorithm on 5-marker windows and explored alternative orders.

In the second step, “comprehensive” maps were obtained in MapMaker by sequentially adding markers from the full set of SNPs that met the more lenient criteria described above to the initial map. Markers were added if they satisfied a log-likelihood threshold of 2.0 for the marker positions, followed by a second round with a log-likelihood threshold of 1.0. We then used the ripple algorithm again on 5-marker windows and explored alternative orders. Typically, MapMaker successfully juxtaposes SNP markers from the same scaffold. However, in marker dense regions with low recombination rates, the likelihoods of alternative marker orders coalesce. In such regions, when multiple markers from the same genomic scaffold were interspersed by markers from a different scaffold, we repositioned the former markers by forcing them in the map together. If the log-likelihood of the map decreased by more than 3.0 (factor 1000), only one of the markers from that scaffold was used in the map. The comprehensive maps provide a balance between marker density and confidence in marker ordering and spacing.

The third step was to create “dense” maps. We added all remaining markers that were not yet incorporated in step two, first at a log-likelihood threshold of 0.5, followed by another round at a log-likelihood threshold of 0.1. We then used the ripple command as described above. The dense maps are useful for anchoring of scaffolds and for obtaining the highest possible resolution of variation in recombination rates, but with the caveat that there is some uncertainty in marker order. Uncertainty is expected to be higher towards the centers of the linkage groups where crossing over events between adjacent markers become substantially less frequent (see Results).

### Comparative analyses

Based on the recent divergence times and high interbreeding successes, we predict a large degree of collinearity of the linkage maps. We note that interpretations must take into account the non-independence of the ParKoh and PruKoh/KonPar maps, as only comparing PruKoh and KonPar comprises a fully independent contrast. We first examined whether inversions or other chromosomal rearrangements were common (affecting linkage map lengths and marker orders) or whether maps were generally collinear by comparing among the initial and comprehensive linkage maps visually using map graphs from MapChart (Voorrips 2002). Inverted or transposed markers present in two or all maps can be detected by connecting “homologs” in MapChart (a homolog in this case means a scaffold that is represented in two or more maps). Then, we tested whether linkage maps are generally collinear across the species pairs quantitatively. We used Spearman’s rank order correlation (ρ) test to examine the strength of correlation between the order in shared markers (the homologs in MapChart). We calculated ρ and the corresponding *P*-value (the probability of observing the measured or stronger correlation given no true correlation exists) by using the cor.test() function in R (R Development Core Team 2016).

We then tested for genetic incompatibilities among the genomes of the four species, by measuring segregation distortion in sliding, 10 cM windows. Although we filtered out markers with very high levels of segregation distortion (using a 5% FDR cutoff) to purge markers with potential sequencing errors, groups of distorted markers in a single region of a linkage group represent genomic regions with biased parental allele contributions, suggesting genetic incompatibilities (or, less common, selfish alleles and other active segregation distorters). Because *L. kohalensis* and *L. paranigra* are more distantly related to each other (and, thus, allowing more time for genetic incompatibilities to accumulate) than they are to *L. pruna* and *L. kona*, respectively (Mendelson and Shaw 2005; see Fig 1.), we expected more regions with significant segregation distortion in the ParKoh map relative to the KonPar and PruKoh maps. We calculated genotype frequency and the negative 10-base logarithm of the *P* value for the *χ* ^2^-square test of deviation from Mendelian inheritance across the linkage groups in R using the R/qtl package (Broman *et al.* 2003). Windows with *P* < 0.01 were considered to have significant segregation distortion, und thus potentially reflecting genetic incompatibilities.

After establishing that the linkage maps were generally collinear (see Results), we merged the maps and examined patterns of variation in crossing over along the *Laupala* genome. Maps were consolidated using ALLMAPS (Tang *et al.* 2015). Then, we calculated species-specific average recombination rates for the linkage groups by dividing the total length of the linkage group (in cM) by the physical length of the pseudomolecule (in million bases, Mb) obtained by merging homologous linkage groups using ALLMAPS. Lastly, to evaluate recombination rate variation along the linkage groups, we fitted smoothing splines (with 10 degrees of freedom, based on the fit of the spline to the observed data) in R to describe the relationship between the consensus physical distance (as per the anchored scaffolds) and the genetic distance specific to each linkage map. Variation in the recombination rate was then assessed by taking the first derivative (i.e. the rate) of the fitted spline function. The estimated recombination rates are likely to be an overestimate of the true recombination rate, because unplaced/unordered parts of the assembly do not contribute to the physical length of the pseudomolecules but are reflected in the genetic distances obtained from crossing-over events in the recombining hybrids.

To test the hypothesis that linked trait and preference genes reside in low recombination regions, we integrated the AFLP map and song and preference QTL peaks identified in previous work on *L. kohalensis* and *L. paranigra* (Shaw and Lesnick 2009) with the current ParKoh SNP map and projected the QTL peaks onto the anchored genome. The SNPs used in the present study were obtained from the same mapping population (same individuals) as in the 2009 AFLP study. Therefore, we combined the high confidence SNPs described above (for the “initial” map) with the AFLP markers reported in (Shaw *et al.* 2007) that were of the same individuals as the SNP markers used in this study and created a new linkage map using the same stringent criteria as for the “initial” maps described above. We projected this map onto the anchored draft genome based on common markers (scaffolds). We then approximated the physical location of the QTL peaks by looking for SNP markers on scaffolds present in the draft genome flanking AFPL markers underneath the QTL peaks identified in the 2009 study.

## DATA ACCESIBILITY

Supplementary files are available on FigShare. See section “supplementary materials” for details. Raw data (vcf files, linkage maps, pseudomolecule agp file), and R-scripts will be deposited on FigShare after final acceptance and are available upon request. The genome assembly and sequencing reads are available on NCBI’s GenBank under BioProject number PRJNA392944. The Genotype-by-sequencing reads will be made available in NCBI’s short read archive under BioProject number PRJNA429815

## RESULTS

### De novo genome assembly

The sequencing of the four libraries yielded 162.5 Gb of raw sequences (Table 1). After read processing, 145.5 Gb was used for the sequence assembly. We compared among assemblies resulting from different Kmer sizes (k = 31, 39, 47, 55, 63, 71, 79 and 87). Based on the N50/L50 and the total assembly size, the assembly produced with k = 87 was retained for the final draft genome. Despite a large number of scaffolds in the final assembly (149,424), the median length of the scaffolds was high and the total length of the assembly covers about 83% of the expected complete genome in *Laupala*.

Gene space coverage in the assembly was evaluated using the *L. kohalensis* cricket gene index (Danley *et al.* 2007) (release 2.0), RNASeq from *Laupala cerasina* (Blankers *et al.* 2018), and by performing a BUSCO search using eukaryotic and arthropod specific conserved genes. Respectively 95% and 92% of the *Laupala* gene index and RNAseq sequences mapped to the current genome. In addition, the BUSCO search indicated very few missing genes in either database (Table 1).

**Table 1.** *Laupala kohalensis* sequencing, assembly and gene space evaluation statistics.

| Sequencing Statistics<br>Library | Raw data |  | Processed data |  |
| --- | --- | --- | --- | --- |
|  | Size (Gb) | Coverage<sup>a</sup> | Size (Gb) | Coverage<sup>a</sup> |
| Paired End 0.2 Kb inserts | 28.9 | 15 | 26.1 | 14 |
| Paired End 0.5 Kb inserts | 63.1 | 33 | 59.8 | 31 |
| Mate Pair 2 Kb inserts | 36.2 | 19 | 31.8 | 17 |
| Mate Pair 5 Kb inserts | 34.3 | 18 | 27.8 | 14 |
| Total | 162.5 | 85 | 145.5 | 76 |

| Assembly Statistics | Contigs | Scaffolds |
| --- | --- | --- |
| Total assembly size (Gb) | 1.6 | 1.6 |
| Total assembled sequences | 219,073 | 148,874 |
| Longest sequence length (Kb) | 465 | 4,541 |
| Average sequence length (Kb) | 7.2 | 10.7 |
| N90 index<sup>b</sup> | 40,926 | 3,505 |
| N90 length (Kb) | 7.7 | 67.7 |
| N50 index | 9,917 | 756 |
| N50 length (Kb) | 43.6 | 583 |
| GC content (%) | 34.9% | 34.9% |

| Gene Space Statistics | Mapping percentage |
| --- | --- |
| *Laupala* unigenes from the Gene Index | 95% |
| *Laupala* RNASeq reads | 92% |

| BUSCO database | Complete | Single copy | Duplicated | Fragmented | Missing | Total |
| --- | --- | --- | --- | --- | --- | --- |
| Eukaryota_odb9 | 98.7% | 93.7% | 5.0% | 0.3% | 1.0% | 303 |
| Arthropoda_odb9 | 99.3% | 96.8% | 2.5% | 0.1% | 0.6% | 1066 |

<sup>a</sup>Coverage is based in an estimated genome size of 1.91 Gb (Petrov *et al.* 2000)
<sup>b</sup>When ordering all contigs (or scaffolds) by size, the N50 or N90 index indicates the number of the longest sequences (contigs or scaffolds) that contain 50% or 90%, respectively, of the total assembled sequence. The N50 and N90 length indicate the length of the shortest sequence in the set of the largest contigs (or scaffolds) that contain 50% or 90%, respectively, of all the sequence in the assembly.

### Collinear linkage maps

We obtained 815,109,126; 522,378,849; and 311,558,401 reads after demultiplexing for ParKoh, KonPar, and PruKoh, respectively. Average sequencing depth ± standard deviation across all individuals in the F2 mapping population after filtering, (before and) after marker selection based on segregation distortion and ancestry information for linkage mapping was (62.4 x ± 162.5) 52.2 x ± 31.4, (44.3 x ± 58.5) 38.1 x ± 23.8, and (56.1 x ± 105.7) 41.8 x ± 29.3, respectively.

In the initial maps, 158 (ParKoh), 170 (KonPar), and 138 (PruKoh) markers were grouped into eight linkage groups at a LOD score of 5.0, corresponding to the seven autosomes and one X-chromosome in *Laupala.* The corresponding marker spacing was 5.14, 4.85, and 7.33 cM. The comprehensive maps contained 526, 650, and 325 markers with an average marker spacing of 1.91, 1.37, and 3.25 cM and on the dense maps we placed 608, 823, and 383 markers with on average 1.69, 1.37, and 3.25 cM. between markers

The recent divergence times and the limited levels of post-zygotic isolation observed in this system led us to hypothesize that the linkage maps would show a high degree of collinearity. The visual comparison of marker positioning showed that the relative locations of shared scaffolds were similar across the linkage maps in both the initial and the comprehensive maps (Fig 2, Fig S1). However, we also observe substantial variation in the total genetic length of homologous linkage groups, indicating recombination rate variation (Fig 2, Fig S1). This variation may in part result from chromosomal rearrangements. However, we can only reliably detect rearrangements in our maps if they are not segregating within species *and* are fixed for alternative arrangements between *L. pruna* and *L. kohalensis* on the one side and *L. paranigra* and *L. kona* on the other side. In that specific scenario the inverted marker order is visible when contrasting the PruKoh and KonPar maps, while the ParKoh map would show reduced recombination in that area (Fig 1C). Despite the apparent variation in recombination rates among homologous linkage groups, Spearman’s rank correlation of pairwise linkage group comparisons was high (ρ varied between 0.91 and 1.00) and similar to values seen in comparisons of intraspecific linkage maps (e.g. Poursarebani *et al.* 2013); the quantitative measure of collinearity was largely consistent across linkage groups and across cross types (Table 2). Finally, merging the maps into a consensus pseudomolecule assembly allowed us to measure the error between individual maps and the merged assembly. Correlations between linkage maps and the pseudomolecule assembly were generally high (> 0.95), indicating substantial synteny (Fig S2).

**Figure 2.**
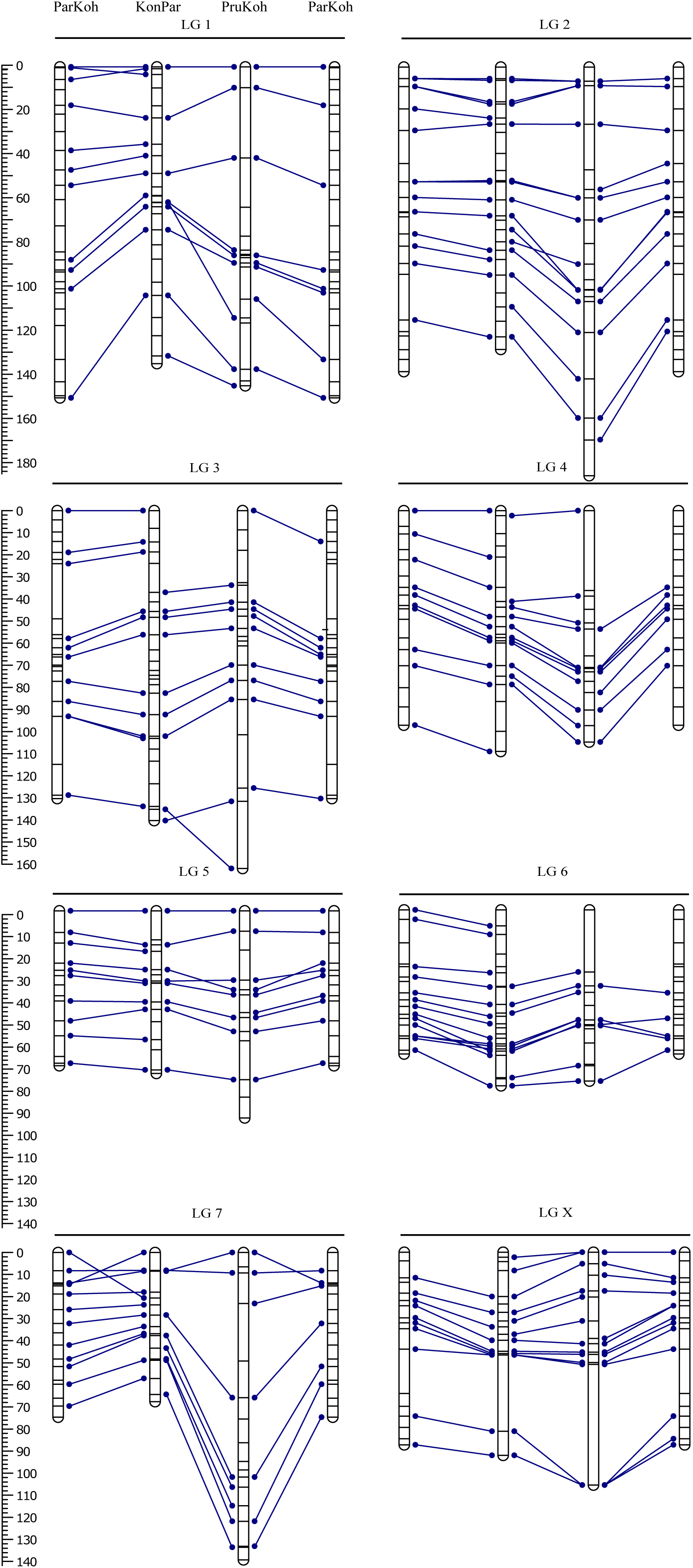
Initial linkage maps. Bars represent linkage groups (LG) for ParKoh, KonPar, and PruKoh. Lines within the bars indicate marker positions. The scale on the left measures marker spacing in cM. Blue lines connect markers on the same scaffold between the different maps. The map for ParKoh is shown twice to facilitate comparison across all three maps. See Fig S1 for comprehensive maps.

**Table 2.** Linkage map comparison. Spearman’s rank correlation (ρ) is shown for each pairwise comparison of linkage maps across all 8 linkage groups.

|  | ParKoh ~ KonPar | ParKoh ~ PruKoh | KonPar ~ PruKoh |
| --- | --- | --- | --- |
| 1 | 0.99‡ | 0.90‡ | 0.97‡ |
| 2 | 0.99‡ | 0.96‡ | 0.93‡ |
| 3 | 1.00‡ | 0.98‡ | 0.95‡ |
| 4 | 0.99‡ | 1.00‡ | 0.97‡ |
| 5 | 0.97‡ | 0.98‡ | 0.95‡ |
| 6 | 0.99‡ | 0.94‡ | 0.94‡ |
| 7 | 0.96‡ | 0.93† | 0.91‡ |
| X | 0.92‡ | 0.96‡ | 0.99‡ |

\* $P < 0.01$ ; † $P < 0.001$ ; ‡ $P < 0.0001$

### Limited heterogeneity in segregation distortion

We expected genetic incompatibilities to be more likely to occur in the ParKoh cross than in the KonPar and PruKoh cross, because *L. kohalensis* and *L. paranigra* are more distantly related than any of the other species pairs (Fig 1). We tested this hypothesis by examining the degree of segregation distortion in markers within 10 cM sliding windows across the linkage maps. Overall, segregation distortion was limited and average genotype frequencies were close to Mendelian expectations (Fig 3). However, LG3 showed a bias against *L. kohalensis* homozygotes in the ParKoh cross but not in any of the other crosses. Additionally, there was significant variation in the frequency of heterozygotes across the linkage groups (linear model Freq[heterozygotes]∼LG x cross: *R*^2^ = 0.21, F_20,1547_ = 20.7, *P* < 0.0001). Post-hoc Tukey Honest Significant Differences revealed that linkage group 7 had the lowest abundance of heterozygotes overall and within each of the intercrosses and that levels of heterozygosity on LG 7 were similar across the maps (Table S2). Together, these results show that from some LGs and in some crosses, certain genotype combinations were less common than expected, potentially as a result for genetic incompatibilities or meiotic drive.

**Figure 3.**
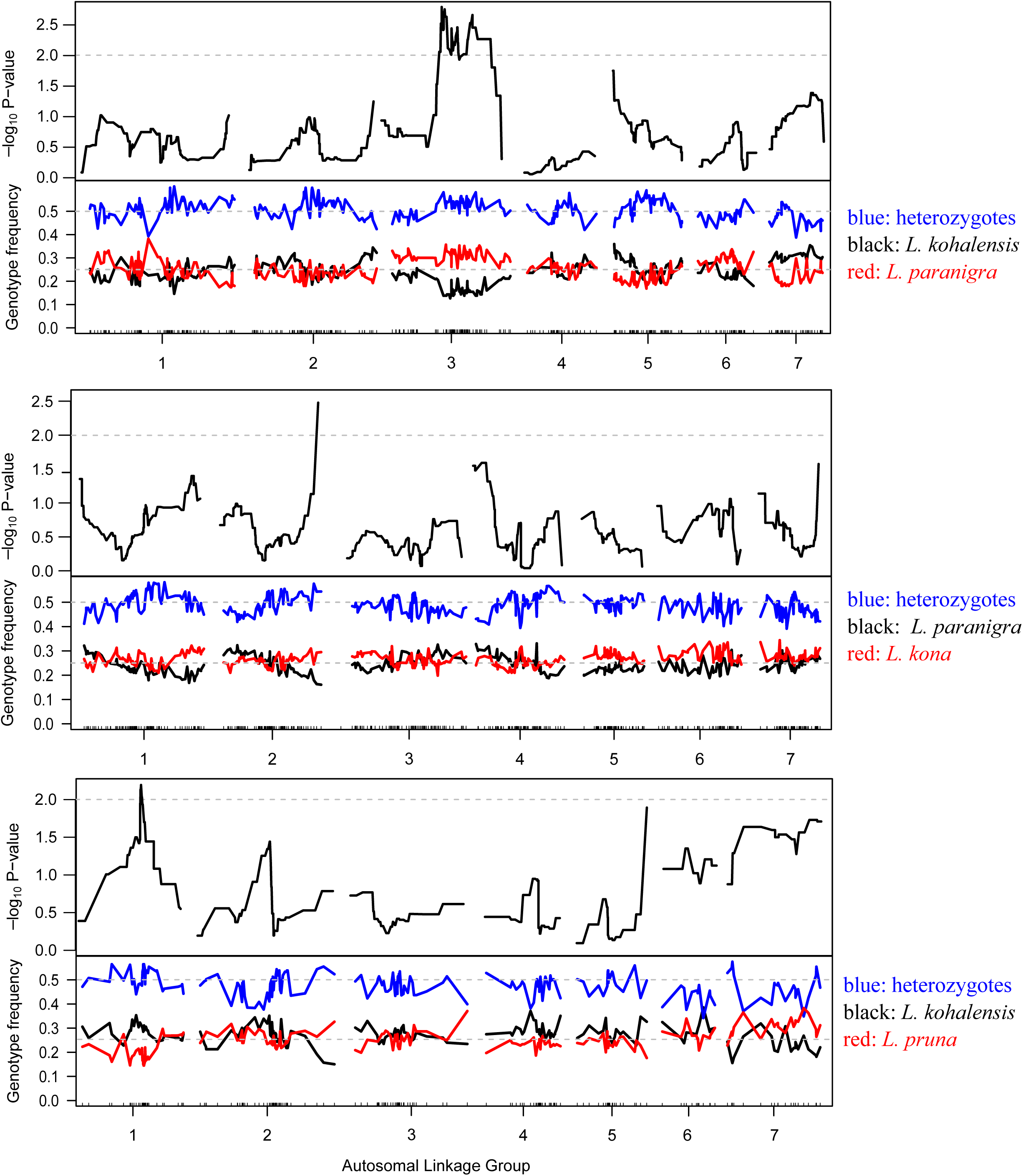
Segregation distortion. For each of the seven autosomal linkage groups within the three comprehensive linkage maps (from top to bottom: ParKoh, KonPar, PruKoh), a sliding window of the negative log-transformed P-values for the χ^2^-square test for deviation from a 1:2:1 segregation ratio is shown across markers with black lines in the top panels. In the panel below, the trace of the frequency of heterozygote genotypes (blue lines) and homozygote genotypes for both parental alleles (black and red lines, respectively) is shown. For each intercross, dashed grey lines indicate P = 0.01 (top panels) or expected allele frequencies based on 1:2:1 inheritance (bottom panels).

### Variable recombination rates across the genome

We anchored a total of 1054 scaffolds covering 720 million base pairs, a little below half the current genome assembly (see Table S3 for scaffold number, N50, and assembly size per LG and Fig S3 for coverage variation across the linkage groups). This gives us enough power to make inferences about broad-scale recombination rate variation, but not about the existence of small-scale recombination hotspots. Average recombination rates (cM/Mb) varied from between 0.75 (KonPar) and 0.93 (ParKoh) on the X chromosome to between 3.12 (KonPar) and 4.24 (PruKoh) on LG6 (Table 3). We note that the recombination rate for LG6 might be artificially inflated because of lower assembly quality (expressed as N50) of this LG relative to the other LGs in all linkage maps and in the pseudochromosomes (Table S3). Both including and excluding the sex chromosome, there is a significant linear relationship between chromosome size and genetic length (linear mixed effect model with cross as random variable; With X: *β* = 0.62, F_1,23_ = 14.95, *P* = 0.0008; without X: *β* = 0.69, F_1,20_ = 29.7, *P* < 0.0001) and between chromosome size and broad-scale recombination rate (with X: *β* = −34.1, F_1,23_ = 29.1, *P* < 0.0001).; without X: *ß* = - 24.0, F1,23 = 63.7, *P* < 0.0001).

**Table 3.**
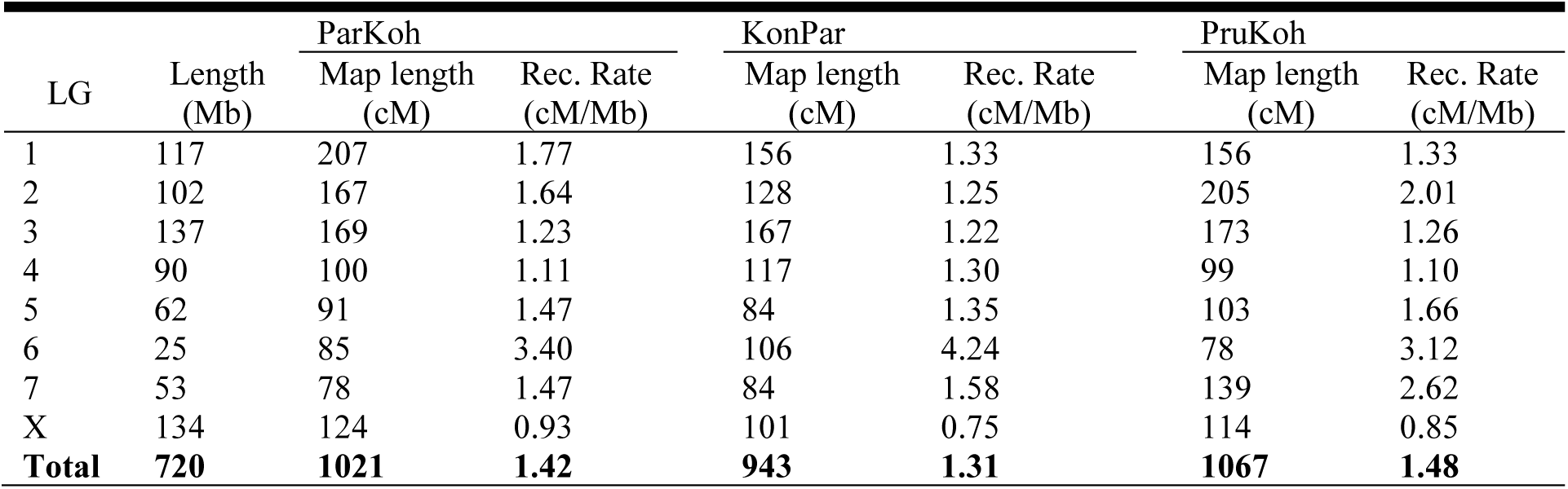
Linkage map summary statistics.

| LG | Length<br>(Mb) | ParKoh |  | KonPar |  | PruKoh |  |
| --- | --- | --- | --- | --- | --- | --- | --- |
|  |  | Map length<br>(cM) | Rec. Rate<br>(cM/Mb) | Map length<br>(cM) | Rec. Rate<br>(cM/Mb) | Map length<br>(cM) | Rec. Rate<br>(cM/Mb) |
| 1 | 117 | 207 | 1.77 | 156 | 1.33 | 156 | 1.33 |
| 2 | 102 | 167 | 1.64 | 128 | 1.25 | 205 | 2.01 |
| 3 | 137 | 169 | 1.23 | 167 | 1.22 | 173 | 1.26 |
| 4 | 90 | 100 | 1.11 | 117 | 1.30 | 99 | 1.10 |
| 5 | 62 | 91 | 1.47 | 84 | 1.35 | 103 | 1.66 |
| 6 | 25 | 85 | 3.40 | 106 | 4.24 | 78 | 3.12 |
| 7 | 53 | 78 | 1.47 | 84 | 1.58 | 139 | 2.62 |
| X | 134 | 124 | 0.93 | 101 | 0.75 | 114 | 0.85 |
| **Total** | **720** | **1021** | **1.42** | **943** | **1.31** | **1067** | **1.48** |

Most linkage groups showed wide regions of strongly reduced recombination rates in the center of the linkage groups (Fig. 4). The general pattern of peripheral peaks in recombination rates juxtaposing large recombination “desserts” was similar among the three intercrosses, but some additional cross-specific peaks in recombination rates were observed on almost all linkage groups (Fig 4).

**Figure 4.**
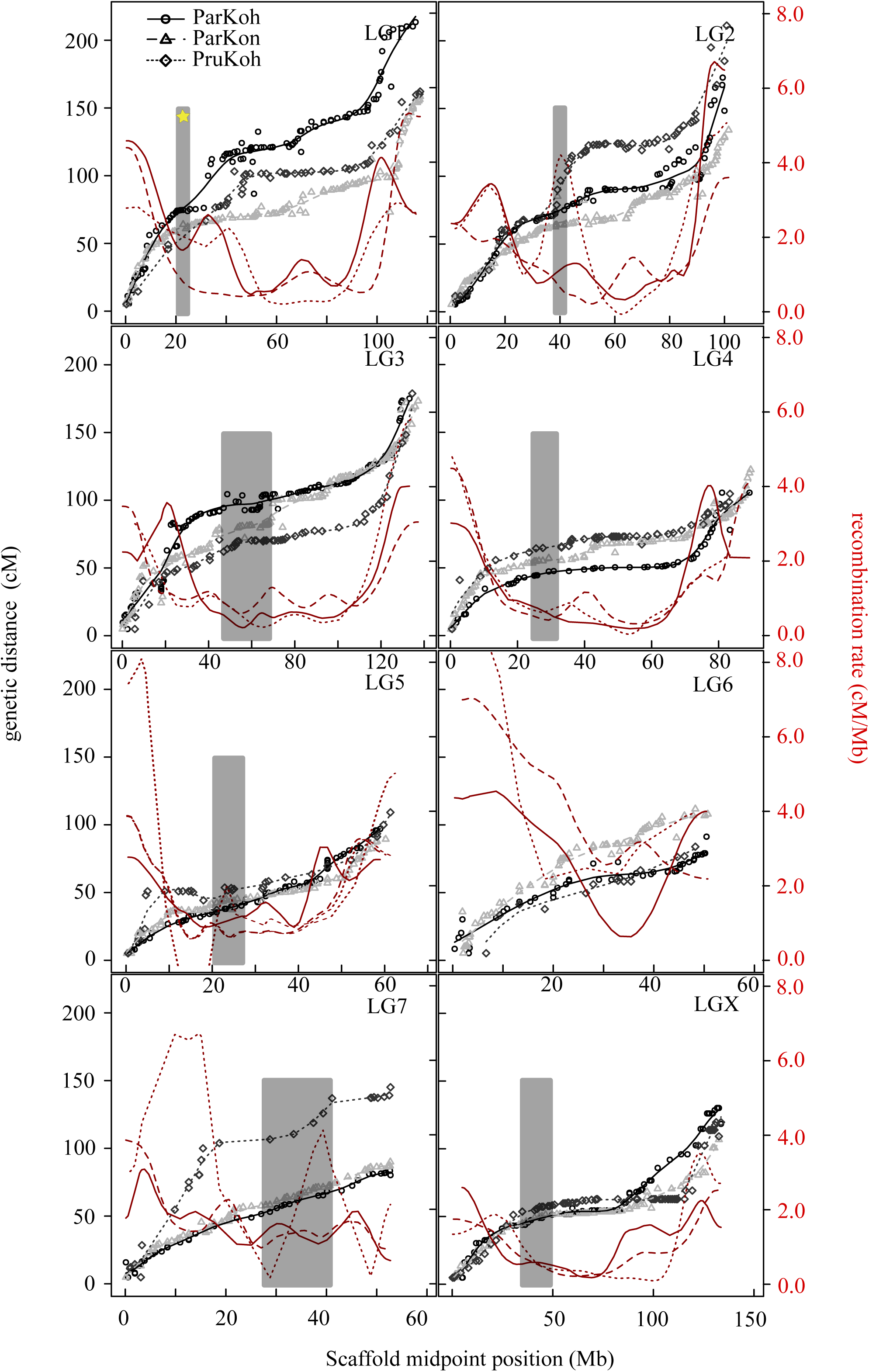
Recombination and Marey maps. Gray-scale symbols and lines indicate the relationship between the physical distance (scaffold midposition) in million base pairs on the x-axis and the genetic distance in cM for each of the 8 linkage groups on the left y-axis. Open dots represent the dense ParKoh linkage map, triangles and diamonds that of the KonPar and PruKoh cross, respectively. The corresponding lines (ParKoh: solid, KonPar: dashed, PruKoh: dotted) indicate the fitted smoothing spline (10 degrees of freedom). The red lines (same stroke style) show the first order derivative of the fitted splines and represent the variation in recombination rate (in cM per Mb, on the right y-axis) as a function of physical distance. Grey bars indicate the approximate location of male song rhythm QTL peaks. The yellow star in the LG1 panel highlights the QTL peak that co-localizes with a female preference QTL peak (Shaw & Lesnick 2009).

### Trait-preference co-evolution despite high recombination

Contrary to our expectation, the approximate location of the colocalizing song and preference QTL peak from (Shaw and Lesnick 2009) was associated with average recombination rates in the ParKoh and KonPar map and low recombination rates in the PruKoh map (Fig 4; Table S4). However, most other QTL peaks are located in regions of low recombination (Fig 4; Table S4).

## DISCUSSION

The evolutionary trajectory of diverging populations and the likelihood of speciation can be heavily influenced by recombination. Within species, recombination can create favorable combinations of alleles or decouple deleterious from beneficial alleles. Among species, regions with low recombination can provide a genetic shield against introgression of maladaptive loci (Noor *et al.* 2001; Rieseberg 2001; Butlin 2005; Slatkin 2008; Noor and Bennett 2009; Cutter and Payseur 2013; Ortiz-Barrientos *et al.* 2016). Understanding recombination is thus critical to understanding adaptation and speciation. Recombination also has important implications for the analysis of genotype-phenotype relationships (Mackay 2001), demographic inference (Li and Durbin 2011), and analyses of genomic variation (Cutter and Payseur 2013; Wolf and Ellegren 2016). However, we still have limited insight into the patterns of recombination rate variation among species and across genomes, in particular for radiations powered largely by behavioral isolation.

Here, we study four species of sexually divergent Hawaiian swordtail crickets and generate the first pseudomolecule-level assembly for Orthoptera and the first published genome assembly for crickets, an important model system in neurobiology, behavioral ecology, and evolutionary genetics (Horch *et al.* 2017). Below, we discuss how our results provide insight into the potential for structural variation (linkage map collinearity) and genetic incompatibilities to drive reproductive isolation among closely related *Laupala* species. We also elaborate on the patterns of variation in recombination rates across the genome. We then discuss the surprising finding that colocalizing male song and female preference QTL did not fall in a region with particularly low recombination. This is important because it challenges the hypothesis that co-evolution of traits and preferences is facilitated by locally reduced recombination between recently diverged populations.

### Collinearity of Genetic Maps

Based on the recent divergence (Mendelson and Shaw 2005) and strong premating isolation of *Laupala* species in the absence of conspicuous morphological and ecological differences (Otte 1994; Shaw 1996; Mendelson and Shaw 2005), we expected limited variation in chromosome structure and few signatures of genetic incompatibility between the species. In line with these expectations, we found that linkage groups are collinear across interspecies crosses (Fig 2). This was true both for comparisons of non-independent species pairs (between ParKoh and the other two crosses), as well as for the independent contrast of the PruKoh map versus the KonPar map. We saw some instances where markers occupied regions that may have been translocated or inverted. However, these instances were rare (Fig 2, Fig S1) and recombination rates were similar among homologous linkage groups across the hybrid families (Fig 4). Moreover, quantitative measures of correlation (Spearman’s rank correlation among maps, Pearson’s correlation coefficient between maps and the pseudomolecule assembly) as well as limited segregation distortion (but see discussion of one exception below) both supported the collinearity hypothesis.

Variation in the organization and structure of chromosomes can contribute to postzygotic reproductive isolation after speciation as well as to the speciation process directly (Noor *et al.* 2001; Rieseberg 2001). We conclude that at least for the *Laupala* group that radiated on the Big Island of Hawaii in the last 500,000 years, structural rearrangements have not played a major role in the evolution of reproductive isolation. This is because, similar to two hybridizing *Heliconius* species (Davey *et al.* 2017), we observed that chromosome-wide recombination rates are relatively conserved and large chromosomal rearrangements are absent. We hypothesize that for *Laupala* on the Big Island premating isolation combined with (partial) geographic separation (i.e. low migration rates) provides a sufficiently strong barrier to gene flow between sister species. Indeed, as has been shown in recent models of the role of inversions in speciation, genomic rearrangements can only invade and spread in diverging populations if levels of gene flow and the contribution of structural variation to isolation (by linking adaptive alleles or incompatibilities) is high relative to the strength of assortative mating (Feder *et al.* 2014; Dagilis and Kirkpatrick 2016).

We acknowledge that the power to detect rearrangements and changes in recombination rates is limited by the resolution of our maps. The average spacing of markers is between 1.37 and 3.25 cM. Thus, the upper limit of the magnitude of intervals within which we can detect rearrangements is on the order of 10^5^ and 10^6^ bp. Due to constraints on the sample size and sequencing strategy, it is thus difficult to attribute subtle variation in marker order and genetic distance between the maps to genomic rearrangements versus mapping errors and sampling variance. Closely related organisms typically show conserved recombination rates within 500 kb intervals; more heterogeneity might be revealed at higher resolution (Stevison *et al.* 2017).

### Genetic incompatibilities

We expected genetic incompatibilities to be more likely between genomes of more distantly related species. Accordingly, we discovered a single region covering approximately half of linkage group 3 with high segregation distortion in ParKoh; we found no such deviations in the other two crosses (Fig 3). Inspection of genotype frequencies indicated that there were fewer individuals than expected that were homozygous for *L. kohalensis* alleles for loci in this region. In a controlled cross, segregation distortion can be caused by prezygotic effects such as meiotic drive of selfish genetic elements and distorter genes (e.g. like *sd* in *Drosophila melanogaster* (Larracuente and Presgraves 2012)), and by postzygotic genetic incompatibilities (Dobzhansky 1937; Muller 1942; Burt and Trivers 2006; Presgraves 2010; Hallmann *et al.* 2017). Genotypic errors may produce superficially similar patterns but are unlikely to distort segregation over large genomic regions and with consistent bias towards the same genotypes. Although meiotic drive is a possible alternative to genetic incompatibilities, we do not see the same effect in the other cross involving *L. paranigra*, where selfish genetic elements or segregation distorters ought to have a similar effect. Overall, the large region on linkage group 3 reveals a potential local post-mating barrier to gene flow that could contribute to strengthening existing prezygotic barriers in secondary contact zones or following episodes of migration.

### Recombination landscape

Chromosomal rearrangements influence genomic divergence by locally altering recombination rates within and among species. Felsenstein (1974, 1981) illuminated the role of *intraspecific* recombination in purging deleterious alleles and the role of *interspecific* recombination in decoupling co-adapted alleles. In recent years, the role of recombination and its interaction with divergent selection and adaptation on genomic scales have received considerable attention (e.g. Yeaman and Whitlock 2011; Feder *et al.* 2012; Samuk *et al.* 2017) and technological advances are shifting focus towards characterizing the recombination landscape across species (Butlin 2005; Slatkin 2008; Noor and Bennett 2009; Barb *et al.* 2014; Burri *et al.* 2015).

Here, we show that there is limited variation in recombination rates across the maps of three interspecific crosses (Fig 4), but strong heterogeneity in recombination rates across the genome. Genome-wide average interspecific recombination rate varied between 1.3 and 1.5 cM/Mb (Table 3), similar to intraspecific rates observed in dipterans and substantially lower than social hymenopterans and lepidopterans (Wilfert *et al.* 2007). We note that our estimates are derived from interspecific maps, which may lead to somewhat lower estimates compared to intraspecific maps (e.g. Beukeboom *et al.* 2010), where genetic incompatibilities and rearrangements may reduce rates of crossing over; however, differences between intra and interspecific recombination might be negligible if rearrangements are rare (e.g., Davey *et al.* 2017). Moreover, we anchored about 50% of the nucleotides in the draft assembly to linkage groups, and there remain many scaffolds not mapped to a genomic position. These ‘missing’ scaffolds are expected to add to the physical length of the chromosomes more so than to the genetic length of the chromosomes, thus lowering the recombination rate. However, our study emphasized relative patterns of recombination, which should not be affected by our sampling. And while we can only approximate intraspecific recombination rates at this point, we note that recent divergence of the species involved and collinearity of the linkage maps support conservation of recombination landscapes across intraspecific and interspecific comparisons.

Interestingly, for all three species pairs we document high variability in interspecific recombination across genomic regions. We found large regions of low recombination in all three maps, with recombination rates well below 1 cM/Mb and occasionally approaching zero, flanked by steep inclines reaching rates up to 6 cM/Mb (Fig 4). This pattern is consistent with earlier findings in plants (Anderson *et al.* 2003), invertebrates (Rockman and Kruglyak 2009; Niehuis *et al.* 2010), and vertebrates (Backström *et al.* 2010; Roesti *et al.* 2013; Singhal *et al.* 2015), but differs from observations in, for example, *Drosophila* (Kulathinal *et al.* 2008) and humans (Myers *et al.* 2005), that show heterogeneity in recombination rates, but not necessarily much higher rates on the periphery of the chromosomes. Commonly invoked drivers of local recombination suppression, such as selection against recombination due to negative epistasis or the maintenance of linkage disequilibrium between mutually beneficial alleles (Smukowski and Noor 2011; Stevison *et al.* 2011; Smukowski Heil *et al.* 2015; Ortiz-Barrientos *et al.* 2016), are not likely to leave chromosome wide signatures. Rather, the observed pattern is more likely attributable to structural properties of chromosomes, such as the location of the centromere and heterochromatin-rich regions (Copenhaver *et al.* 1999; Haupt *et al.* 2001). Roesti *et al*. (2013) observe similar recombination landscapes in stickleback and suggest it might be due to peripheral clustering during meiosis prophase I to facilitate homolog pairing (Harper *et al.* 2004; Brown *et al.* 2005). Regardless of the mechanism, the observed genomic architecture will drive substantial heterogeneity in the propensity of favorable and/or maladaptive alleles to come together, break apart, and introgress in heterospecific backgrounds.

### Trait-Preference Co-evolution

On way in which recombination heterogeneity may be important in the study system is in facilitating trait-preference co-evolution. If trait and preference genes are coupled through physical linkage (Kirkpatrick and Hall 2004), linkage can be stronger and span wider physical distances in regions with reduced recombination. We hypothesized that recombination facilitates linkage between trait and preference genes in *Laupala* because a previous study showed that a major song QTL (∼ 9% of the parental difference in male song) co-localizes with a preference QTL (∼14% of parental difference for female preference) in a cross between *L. kohalensis* and *L. paranigra* (Shaw and Lesnick 2009). Contrary to our expectation, we show that the co-localizing QTL fall in a region with intermediate to high recombination rates (> 2.0 cM) compared to chromosomal averages (typically 1 - 2 cM). This suggests that reduced recombination over larger physical distances is unlikely to be driving trait-preference co-evolution in this system. Importantly, a high speciation rate and wide-spread divergence in sexual signaling phenotypes suggest a primary role for trait-preference co-evolution in *Laupala* speciation (Mendelson and Shaw 2005; Shaw *et al.* 2011). Additionally, although these species have likely diverged in allopatry (Mendelson and Shaw 2005), some level of interspecific gene exchange is likely given historical biogeography, widespread secondary contact and evidence derived from discordant nuclear and mitochondrial gene trees (Shaw 2002).

How then is linkage disequilibrium between traits and preferences maintained? First, QTL may co-localize due to very tight physical linkage or pleiotropy instead of looser linkage. Under these two mechanisms, a lack of physical space for crossing over to occur rather than low recombination rates maintains linkage disequilibrium. Linkage disequilibrium might also persist in the face of recombination if strong assortative mating results from female mate preference. In this case, genetic correlations between the sexes will evolve, coupling signal and preference independent of their genetic distance (Fisher 1930; Lande 1981). Recent simulation studies showed that the probability with which recombination rate modifiers that link co-adaptive alleles spread in a populations is lower when assortative mating is strong, recombination between loci is low, and selection on the loci themselves is strong (Feder *et al.* 2014; Dagilis and Kirkpatrick 2016). Third, the current test involves only a single locus and additional tests are required to more robustly examine the relationship between recombination and trait-preference co-evolution. We observed that several known male song QTL on other linkage groups fall in regions of low recombination. Additional female preference QTL covary with these song QTL as well (Wiley *et al.* 2012) although precise map locations are not yet known.

In summary, we find limited variation in chromosome structure among species, but strong heterogeneity in the recombination landscape across the genome. We present a *de novo* genome assembly and anchor a substantial part of the *L. kohalensis* genome to pseudomolecules. Crickets are an important model system for evolutionary and neurobiological research (Horch *et al.* 2017). but limited genomic resources are available. The first Orthopteran pseudomolecule-level draft genome and recombination rate map are thus important new contributions to future speciation genomics research. This study further provides important insight into the extent to which structural variation and genetic incompatibilities contribute to isolation among closely related, sexually divergent species. We also shine light on the role of recombination in trait-preference co-evolution and argue that current evidence supports that, at least in *Laupala*, the evolution of behavioral isolation is not contingent on structural genomic variation and locally reduced recombination.

## ACKNOWLEDGEMENTS

We thank Stephen Chenoweth and two anonymous reviewers for helpful comments that strongly improved the quality of this manuscript. We further thank the Shaw lab, in particular Mingzi Xu, as well as Michael Sheehan and other members from Cornell’s Neurobiology and Behavior department for input that contributed to the interpretation of the findings. This work was supported by the National Science Foundation (DEB 1241060, IOS 1257682 and IOS 0843528).

## SUPPLEMENTARY MATERIAL

Table S1. Geographic locations of sampled populations

Table S2. Segregation distortion (count of heterozygotes per genotype) statistics. Table S3. Summary statistics for anchored assembly

Table S4. Integrated AFLP and SNP map for the *L. kohalensis* x *L. paranigra* cross

Figure S1. Comprehensive linkage maps.

Figure S2. ALLMAPS output

Figure S3. Coverage per cross per linkage group

**Table S1.** Geographic locations of sampled populations

| Species | Locality Name | Latitude (N) | Longitude (W) |
| --- | --- | --- | --- |
| *L. kona* | Manuka | 19 deg 12' | 155 deg 81' |
| *L. paranigra* | Kaiwiki | 19 deg 46' | 155 deg 10' |
| *L. kohalensis* | Pololu Valley | 20 deg 10' | 155 deg 46' |
| *L. pruna* | Kaiholena | 19 deg 10' | 155 deg 35' |

**Table S2.**
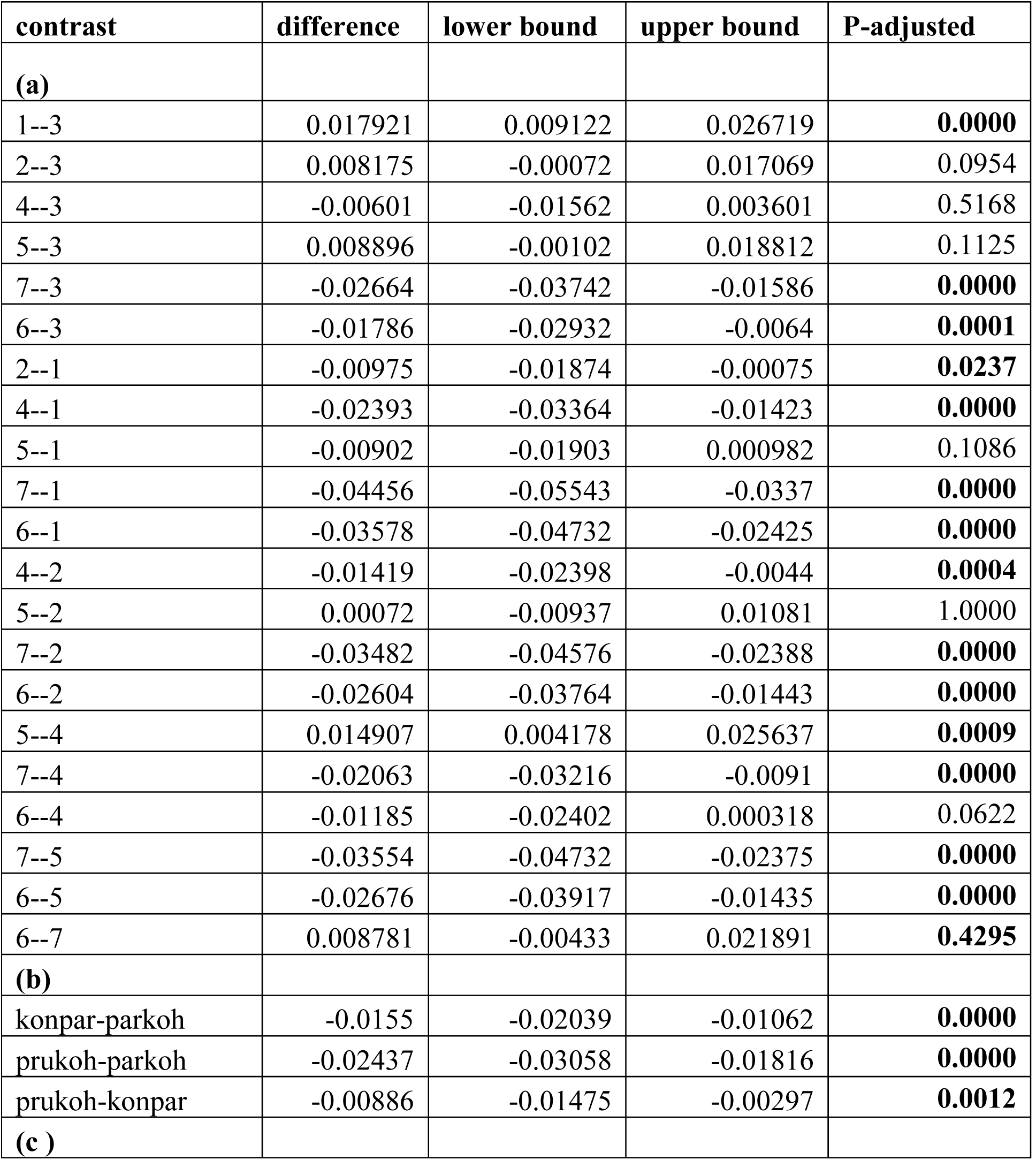
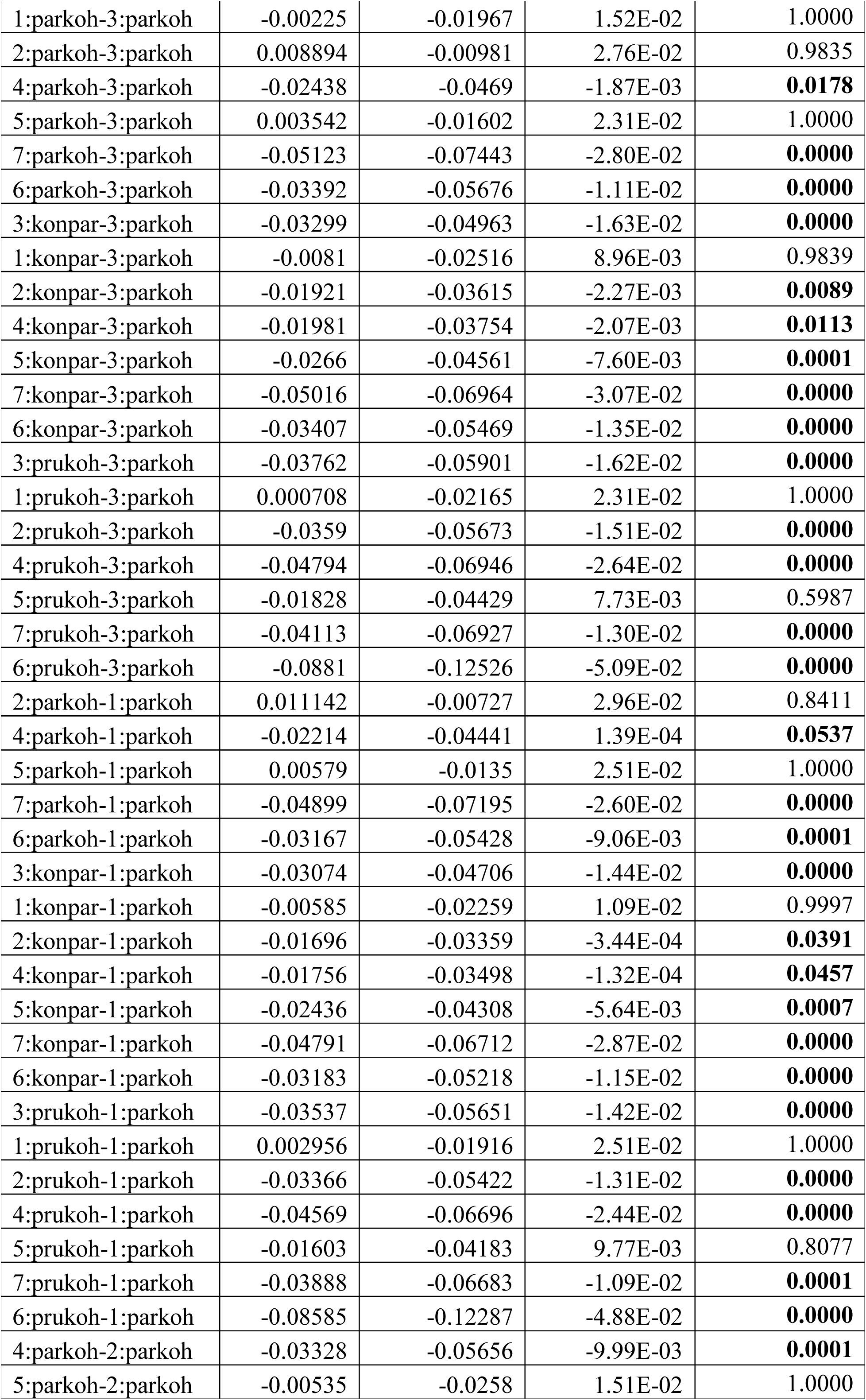
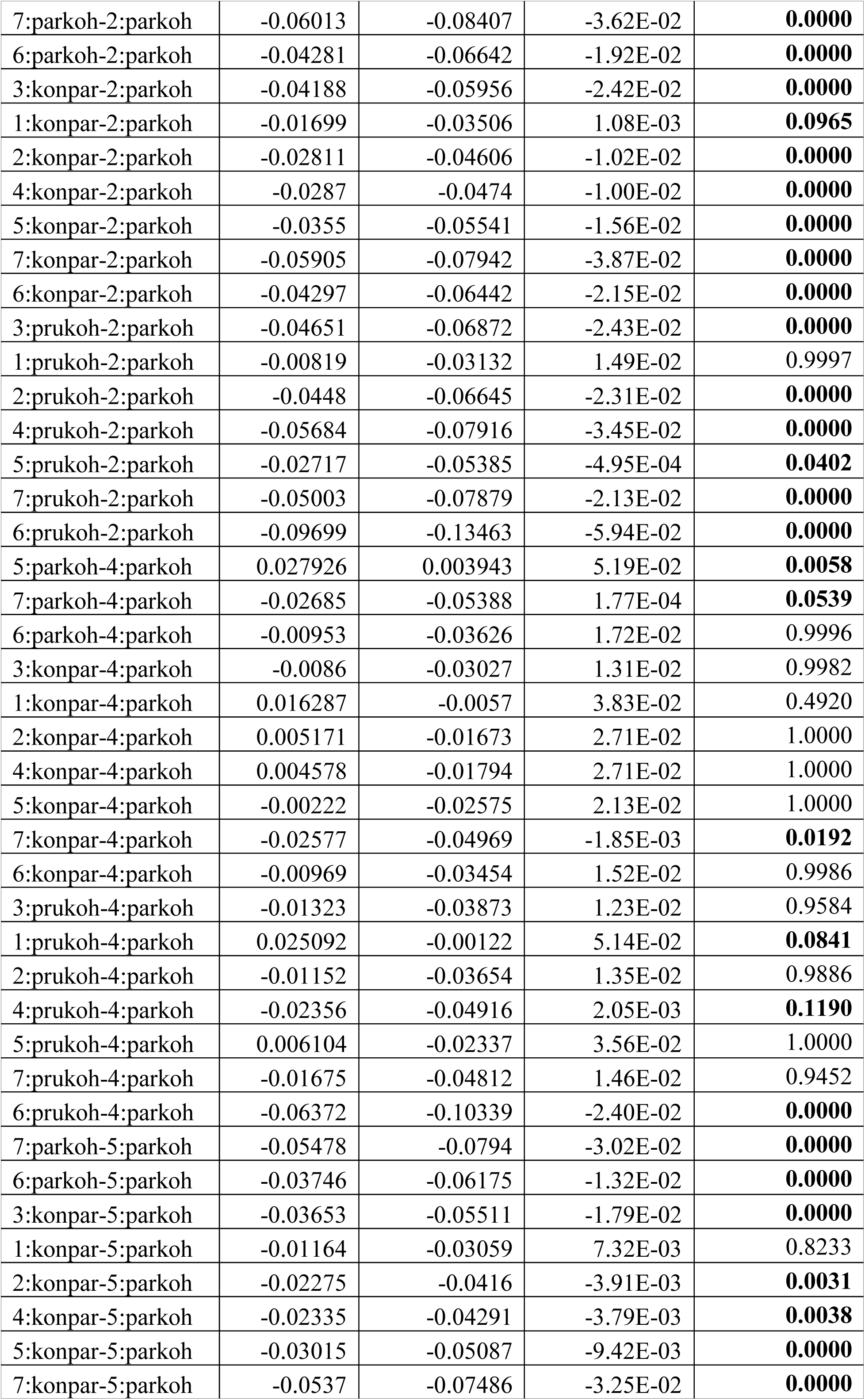
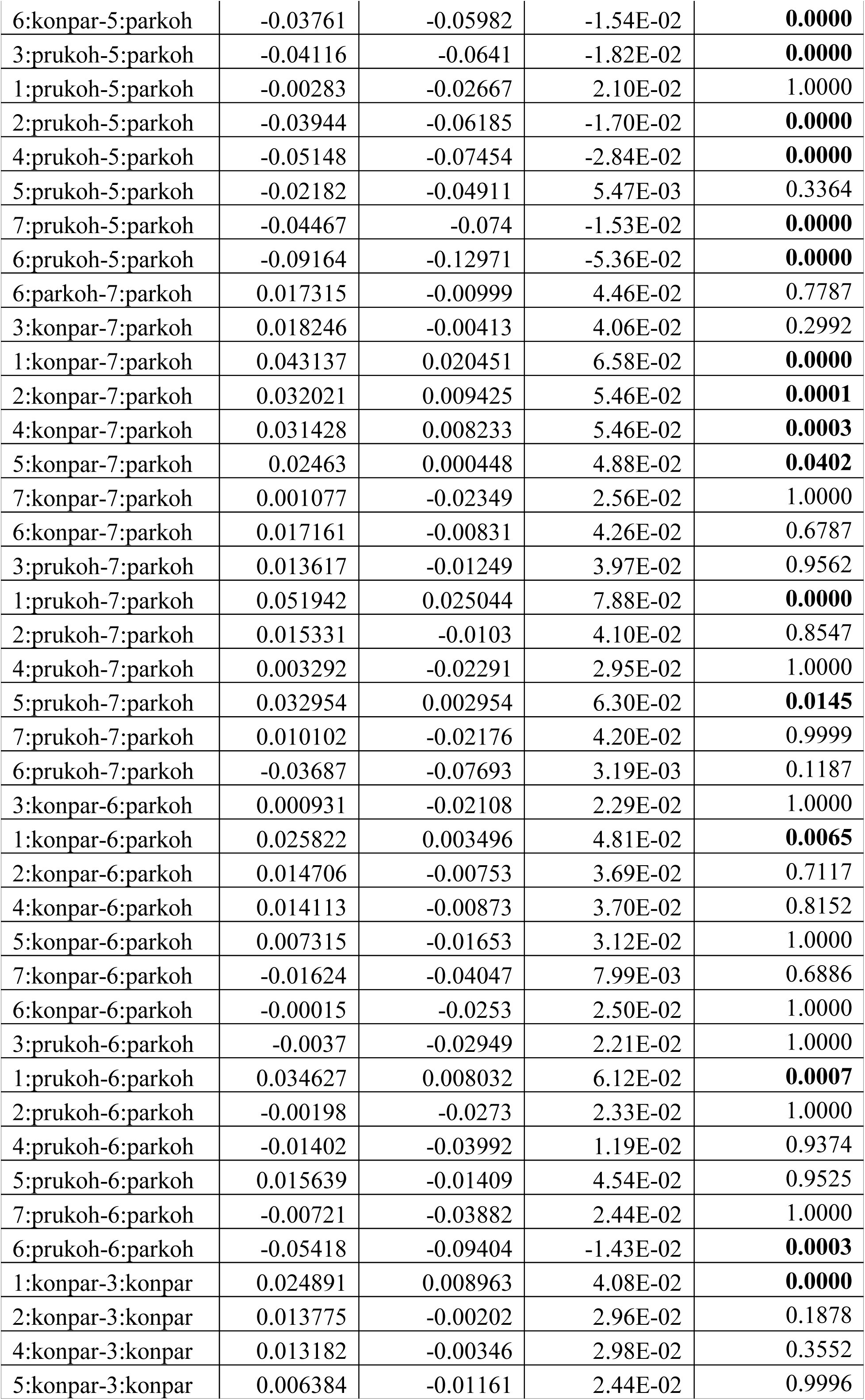
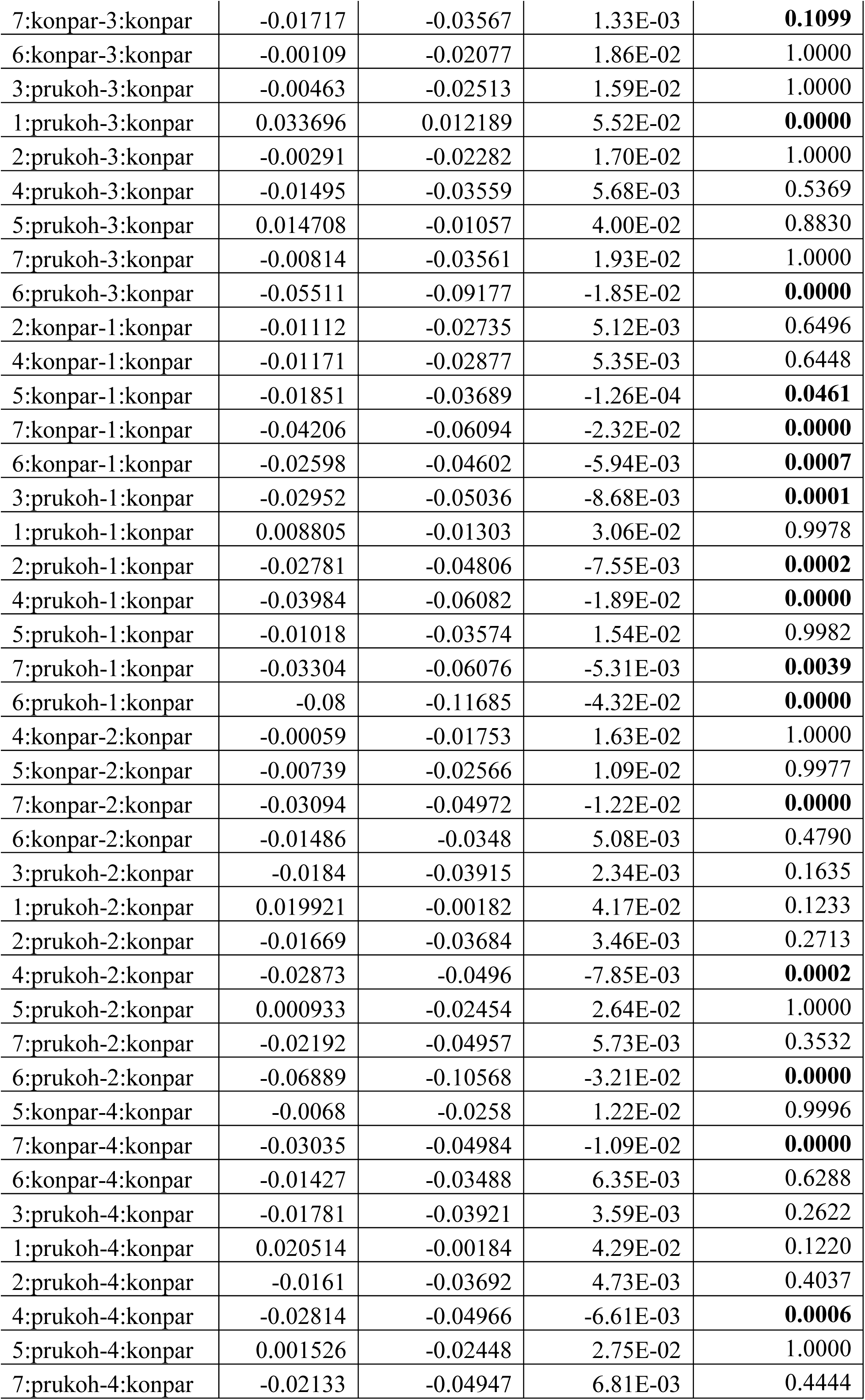
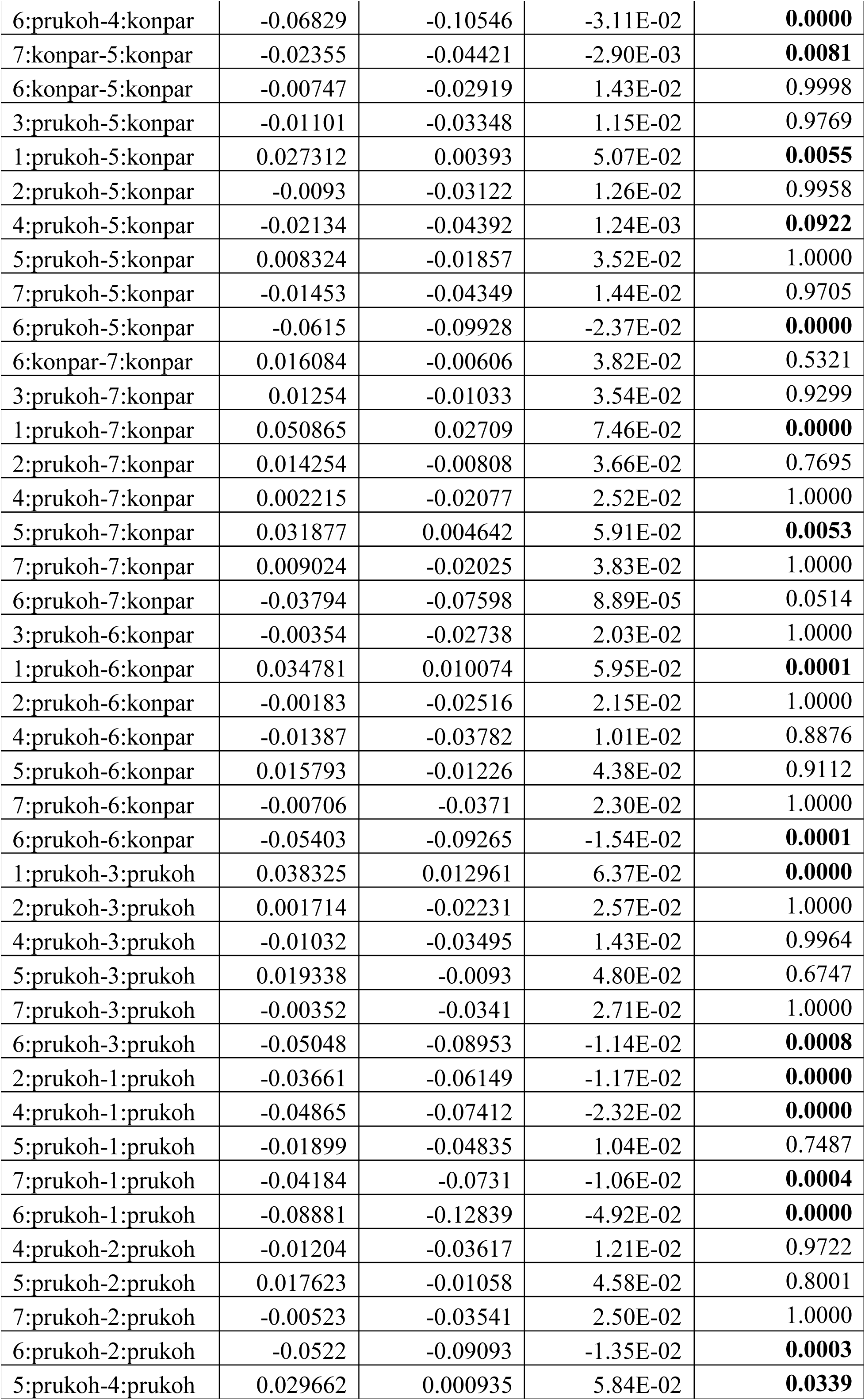
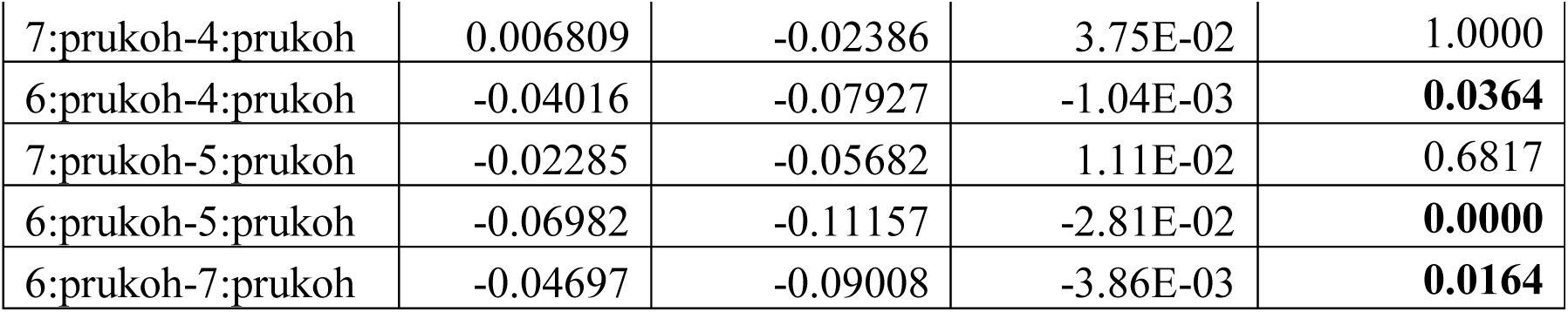
Segregation distortion (count of heterozygotes per genotype) statistics. *post hoc* Tukey Honest Significant Differences corresponding to the number of heterozygotes (a) across linkage groups (b) across species, (c) and across species nested in linkage groups

| contrast | difference | lower bound | upper bound | P-adjusted |
| --- | --- | --- | --- | --- |
| **(a)** |  |  |  |  |
| 1--3 | 0.017921 | 0.009122 | 0.026719 | **0.0000** |
| 2--3 | 0.008175 | -0.00072 | 0.017069 | 0.0954 |
| 4--3 | -0.00601 | -0.01562 | 0.003601 | 0.5168 |
| 5--3 | 0.008896 | -0.00102 | 0.018812 | 0.1125 |
| 7--3 | -0.02664 | -0.03742 | -0.01586 | **0.0000** |
| 6--3 | -0.01786 | -0.02932 | -0.0064 | **0.0001** |
| 2--1 | -0.00975 | -0.01874 | -0.00075 | **0.0237** |
| 4--1 | -0.02393 | -0.03364 | -0.01423 | **0.0000** |
| 5--1 | -0.00902 | -0.01903 | 0.000982 | 0.1086 |
| 7--1 | -0.04456 | -0.05543 | -0.0337 | **0.0000** |
| 6--1 | -0.03578 | -0.04732 | -0.02425 | **0.0000** |
| 4--2 | -0.01419 | -0.02398 | -0.0044 | **0.0004** |
| 5--2 | 0.00072 | -0.00937 | 0.01081 | 1.0000 |
| 7--2 | -0.03482 | -0.04576 | -0.02388 | **0.0000** |
| 6--2 | -0.02604 | -0.03764 | -0.01443 | **0.0000** |
| 5--4 | 0.014907 | 0.004178 | 0.025637 | **0.0009** |
| 7--4 | -0.02063 | -0.03216 | -0.0091 | **0.0000** |
| 6--4 | -0.01185 | -0.02402 | 0.000318 | 0.0622 |
| 7--5 | -0.03554 | -0.04732 | -0.02375 | **0.0000** |
| 6--5 | -0.02676 | -0.03917 | -0.01435 | **0.0000** |
| 6--7 | 0.008781 | -0.00433 | 0.021891 | **0.4295** |
| **(b)** |  |  |  |  |
| konpar-parkoh | -0.0155 | -0.02039 | -0.01062 | **0.0000** |
| prukoh-parkoh | -0.02437 | -0.03058 | -0.01816 | **0.0000** |
| prukoh-konpar | -0.00886 | -0.01475 | -0.00297 | **0.0012** |
| **(c)** |  |  |  |  |
| 1:parkoh-3:parkoh | -0.00225 | -0.01967 | 1.52E-02 | 1.0000 |
| 2:parkoh-3:parkoh | 0.008894 | -0.00981 | 2.76E-02 | 0.9835 |
| 4:parkoh-3:parkoh | -0.02438 | -0.0469 | -1.87E-03 | **0.0178** |
| 5:parkoh-3:parkoh | 0.003542 | -0.01602 | 2.31E-02 | 1.0000 |
| 7:parkoh-3:parkoh | -0.05123 | -0.07443 | -2.80E-02 | **0.0000** |
| 6:parkoh-3:parkoh | -0.03392 | -0.05676 | -1.11E-02 | **0.0000** |
| 3:konpar-3:parkoh | -0.03299 | -0.04963 | -1.63E-02 | **0.0000** |
| 1:konpar-3:parkoh | -0.0081 | -0.02516 | 8.96E-03 | 0.9839 |
| 2:konpar-3:parkoh | -0.01921 | -0.03615 | -2.27E-03 | **0.0089** |
| 4:konpar-3:parkoh | -0.01981 | -0.03754 | -2.07E-03 | **0.0113** |
| 5:konpar-3:parkoh | -0.0266 | -0.04561 | -7.60E-03 | **0.0001** |
| 7:konpar-3:parkoh | -0.05016 | -0.06964 | -3.07E-02 | **0.0000** |
| 6:konpar-3:parkoh | -0.03407 | -0.05469 | -1.35E-02 | **0.0000** |
| 3:prukoh-3:parkoh | -0.03762 | -0.05901 | -1.62E-02 | **0.0000** |
| 1:prukoh-3:parkoh | 0.000708 | -0.02165 | 2.31E-02 | 1.0000 |
| 2:prukoh-3:parkoh | -0.0359 | -0.05673 | -1.51E-02 | **0.0000** |
| 4:prukoh-3:parkoh | -0.04794 | -0.06946 | -2.64E-02 | **0.0000** |
| 5:prukoh-3:parkoh | -0.01828 | -0.04429 | 7.73E-03 | 0.5987 |
| 7:prukoh-3:parkoh | -0.04113 | -0.06927 | -1.30E-02 | **0.0000** |
| 6:prukoh-3:parkoh | -0.0881 | -0.12526 | -5.09E-02 | **0.0000** |
| 2:parkoh-1:parkoh | 0.011142 | -0.00727 | 2.96E-02 | 0.8411 |
| 4:parkoh-1:parkoh | -0.02214 | -0.04441 | 1.39E-04 | **0.0537** |
| 5:parkoh-1:parkoh | 0.00579 | -0.0135 | 2.51E-02 | 1.0000 |
| 7:parkoh-1:parkoh | -0.04899 | -0.07195 | -2.60E-02 | **0.0000** |
| 6:parkoh-1:parkoh | -0.03167 | -0.05428 | -9.06E-03 | **0.0001** |
| 3:konpar-1:parkoh | -0.03074 | -0.04706 | -1.44E-02 | **0.0000** |
| 1:konpar-1:parkoh | -0.00585 | -0.02259 | 1.09E-02 | 0.9997 |
| 2:konpar-1:parkoh | -0.01696 | -0.03359 | -3.44E-04 | **0.0391** |
| 4:konpar-1:parkoh | -0.01756 | -0.03498 | -1.32E-04 | **0.0457** |
| 5:konpar-1:parkoh | -0.02436 | -0.04308 | -5.64E-03 | **0.0007** |
| 7:konpar-1:parkoh | -0.04791 | -0.06712 | -2.87E-02 | **0.0000** |
| 6:konpar-1:parkoh | -0.03183 | -0.05218 | -1.15E-02 | **0.0000** |
| 3:prukoh-1:parkoh | -0.03537 | -0.05651 | -1.42E-02 | **0.0000** |
| 1:prukoh-1:parkoh | 0.002956 | -0.01916 | 2.51E-02 | 1.0000 |
| 2:prukoh-1:parkoh | -0.03366 | -0.05422 | -1.31E-02 | **0.0000** |
| 4:prukoh-1:parkoh | -0.04569 | -0.06696 | -2.44E-02 | **0.0000** |
| 5:prukoh-1:parkoh | -0.01603 | -0.04183 | 9.77E-03 | 0.8077 |
| 7:prukoh-1:parkoh | -0.03888 | -0.06683 | -1.09E-02 | **0.0001** |
| 6:prukoh-1:parkoh | -0.08585 | -0.12287 | -4.88E-02 | **0.0000** |
| 4:parkoh-2:parkoh | -0.03328 | -0.05656 | -9.99E-03 | **0.0001** |
| 5:parkoh-2:parkoh | -0.00535 | -0.0258 | 1.51E-02 | 1.0000 |
| 7:parkoh-2:parkoh | -0.06013 | -0.08407 | -3.62E-02 | **0.0000** |
| 6:parkoh-2:parkoh | -0.04281 | -0.06642 | -1.92E-02 | **0.0000** |
| 3:konpar-2:parkoh | -0.04188 | -0.05956 | -2.42E-02 | **0.0000** |
| 1:konpar-2:parkoh | -0.01699 | -0.03506 | 1.08E-03 | **0.0965** |
| 2:konpar-2:parkoh | -0.02811 | -0.04606 | -1.02E-02 | **0.0000** |
| 4:konpar-2:parkoh | -0.0287 | -0.0474 | -1.00E-02 | **0.0000** |
| 5:konpar-2:parkoh | -0.0355 | -0.05541 | -1.56E-02 | **0.0000** |
| 7:konpar-2:parkoh | -0.05905 | -0.07942 | -3.87E-02 | **0.0000** |
| 6:konpar-2:parkoh | -0.04297 | -0.06442 | -2.15E-02 | **0.0000** |
| 3:prukoh-2:parkoh | -0.04651 | -0.06872 | -2.43E-02 | **0.0000** |
| 1:prukoh-2:parkoh | -0.00819 | -0.03132 | 1.49E-02 | 0.9997 |
| 2:prukoh-2:parkoh | -0.0448 | -0.06645 | -2.31E-02 | **0.0000** |
| 4:prukoh-2:parkoh | -0.05684 | -0.07916 | -3.45E-02 | **0.0000** |
| 5:prukoh-2:parkoh | -0.02717 | -0.05385 | -4.95E-04 | **0.0402** |
| 7:prukoh-2:parkoh | -0.05003 | -0.07879 | -2.13E-02 | **0.0000** |
| 6:prukoh-2:parkoh | -0.09699 | -0.13463 | -5.94E-02 | **0.0000** |
| 5:parkoh-4:parkoh | 0.027926 | 0.003943 | 5.19E-02 | **0.0058** |
| 7:parkoh-4:parkoh | -0.02685 | -0.05388 | 1.77E-04 | **0.0539** |
| 6:parkoh-4:parkoh | -0.00953 | -0.03626 | 1.72E-02 | 0.9996 |
| 3:konpar-4:parkoh | -0.0086 | -0.03027 | 1.31E-02 | 0.9982 |
| 1:konpar-4:parkoh | 0.016287 | -0.0057 | 3.83E-02 | 0.4920 |
| 2:konpar-4:parkoh | 0.005171 | -0.01673 | 2.71E-02 | 1.0000 |
| 4:konpar-4:parkoh | 0.004578 | -0.01794 | 2.71E-02 | 1.0000 |
| 5:konpar-4:parkoh | -0.00222 | -0.02575 | 2.13E-02 | 1.0000 |
| 7:konpar-4:parkoh | -0.02577 | -0.04969 | -1.85E-03 | **0.0192** |
| 6:konpar-4:parkoh | -0.00969 | -0.03454 | 1.52E-02 | 0.9986 |
| 3:prukoh-4:parkoh | -0.01323 | -0.03873 | 1.23E-02 | 0.9584 |
| 1:prukoh-4:parkoh | 0.025092 | -0.00122 | 5.14E-02 | **0.0841** |
| 2:prukoh-4:parkoh | -0.01152 | -0.03654 | 1.35E-02 | 0.9886 |
| 4:prukoh-4:parkoh | -0.02356 | -0.04916 | 2.05E-03 | **0.1190** |
| 5:prukoh-4:parkoh | 0.006104 | -0.02337 | 3.56E-02 | 1.0000 |
| 7:prukoh-4:parkoh | -0.01675 | -0.04812 | 1.46E-02 | 0.9452 |
| 6:prukoh-4:parkoh | -0.06372 | -0.10339 | -2.40E-02 | **0.0000** |
| 7:parkoh-5:parkoh | -0.05478 | -0.0794 | -3.02E-02 | **0.0000** |
| 6:parkoh-5:parkoh | -0.03746 | -0.06175 | -1.32E-02 | **0.0000** |
| 3:konpar-5:parkoh | -0.03653 | -0.05511 | -1.79E-02 | **0.0000** |
| 1:konpar-5:parkoh | -0.01164 | -0.03059 | 7.32E-03 | 0.8233 |
| 2:konpar-5:parkoh | -0.02275 | -0.0416 | -3.91E-03 | **0.0031** |
| 4:konpar-5:parkoh | -0.02335 | -0.04291 | -3.79E-03 | **0.0038** |
| 5:konpar-5:parkoh | -0.03015 | -0.05087 | -9.42E-03 | **0.0000** |
| 7:konpar-5:parkoh | -0.0537 | -0.07486 | -3.25E-02 | **0.0000** |
| 6:konpar-5:parkoh | -0.03761 | -0.05982 | -1.54E-02 | **0.0000** |
| 3:prukoh-5:parkoh | -0.04116 | -0.0641 | -1.82E-02 | **0.0000** |
| 1:prukoh-5:parkoh | -0.00283 | -0.02667 | 2.10E-02 | 1.0000 |
| 2:prukoh-5:parkoh | -0.03944 | -0.06185 | -1.70E-02 | **0.0000** |
| 4:prukoh-5:parkoh | -0.05148 | -0.07454 | -2.84E-02 | **0.0000** |
| 5:prukoh-5:parkoh | -0.02182 | -0.04911 | 5.47E-03 | 0.3364 |
| 7:prukoh-5:parkoh | -0.04467 | -0.074 | -1.53E-02 | **0.0000** |
| 6:prukoh-5:parkoh | -0.09164 | -0.12971 | -5.36E-02 | **0.0000** |
| 6:parkoh-7:parkoh | 0.017315 | -0.00999 | 4.46E-02 | 0.7787 |
| 3:konpar-7:parkoh | 0.018246 | -0.00413 | 4.06E-02 | 0.2992 |
| 1:konpar-7:parkoh | 0.043137 | 0.020451 | 6.58E-02 | **0.0000** |
| 2:konpar-7:parkoh | 0.032021 | 0.009425 | 5.46E-02 | **0.0001** |
| 4:konpar-7:parkoh | 0.031428 | 0.008233 | 5.46E-02 | **0.0003** |
| 5:konpar-7:parkoh | 0.02463 | 0.000448 | 4.88E-02 | **0.0402** |
| 7:konpar-7:parkoh | 0.001077 | -0.02349 | 2.56E-02 | 1.0000 |
| 6:konpar-7:parkoh | 0.017161 | -0.00831 | 4.26E-02 | 0.6787 |
| 3:prukoh-7:parkoh | 0.013617 | -0.01249 | 3.97E-02 | 0.9562 |
| 1:prukoh-7:parkoh | 0.051942 | 0.025044 | 7.88E-02 | **0.0000** |
| 2:prukoh-7:parkoh | 0.015331 | -0.0103 | 4.10E-02 | 0.8547 |
| 4:prukoh-7:parkoh | 0.003292 | -0.02291 | 2.95E-02 | 1.0000 |
| 5:prukoh-7:parkoh | 0.032954 | 0.002954 | 6.30E-02 | **0.0145** |
| 7:prukoh-7:parkoh | 0.010102 | -0.02176 | 4.20E-02 | 0.9999 |
| 6:prukoh-7:parkoh | -0.03687 | -0.07693 | 3.19E-03 | 0.1187 |
| 3:konpar-6:parkoh | 0.000931 | -0.02108 | 2.29E-02 | 1.0000 |
| 1:konpar-6:parkoh | 0.025822 | 0.003496 | 4.81E-02 | **0.0065** |
| 2:konpar-6:parkoh | 0.014706 | -0.00753 | 3.69E-02 | 0.7117 |
| 4:konpar-6:parkoh | 0.014113 | -0.00873 | 3.70E-02 | 0.8152 |
| 5:konpar-6:parkoh | 0.007315 | -0.01653 | 3.12E-02 | 1.0000 |
| 7:konpar-6:parkoh | -0.01624 | -0.04047 | 7.99E-03 | 0.6886 |
| 6:konpar-6:parkoh | -0.00015 | -0.0253 | 2.50E-02 | 1.0000 |
| 3:prukoh-6:parkoh | -0.0037 | -0.02949 | 2.21E-02 | 1.0000 |
| 1:prukoh-6:parkoh | 0.034627 | 0.008032 | 6.12E-02 | **0.0007** |
| 2:prukoh-6:parkoh | -0.00198 | -0.0273 | 2.33E-02 | 1.0000 |
| 4:prukoh-6:parkoh | -0.01402 | -0.03992 | 1.19E-02 | 0.9374 |
| 5:prukoh-6:parkoh | 0.015639 | -0.01409 | 4.54E-02 | 0.9525 |
| 7:prukoh-6:parkoh | -0.00721 | -0.03882 | 2.44E-02 | 1.0000 |
| 6:prukoh-6:parkoh | -0.05418 | -0.09404 | -1.43E-02 | **0.0003** |
| 1:konpar-3:konpar | 0.024891 | 0.008963 | 4.08E-02 | **0.0000** |
| 2:konpar-3:konpar | 0.013775 | -0.00202 | 2.96E-02 | 0.1878 |
| 4:konpar-3:konpar | 0.013182 | -0.00346 | 2.98E-02 | 0.3552 |
| 5:konpar-3:konpar | 0.006384 | -0.01161 | 2.44E-02 | 0.9996 |
| 7:konpar-3:konpar | -0.01717 | -0.03567 | 1.33E-03 | **0.1099** |
| 6:konpar-3:konpar | -0.00109 | -0.02077 | 1.86E-02 | 1.0000 |
| 3:prukoh-3:konpar | -0.00463 | -0.02513 | 1.59E-02 | 1.0000 |
| 1:prukoh-3:konpar | 0.033696 | 0.012189 | 5.52E-02 | **0.0000** |
| 2:prukoh-3:konpar | -0.00291 | -0.02282 | 1.70E-02 | 1.0000 |
| 4:prukoh-3:konpar | -0.01495 | -0.03559 | 5.68E-03 | 0.5369 |
| 5:prukoh-3:konpar | 0.014708 | -0.01057 | 4.00E-02 | 0.8830 |
| 7:prukoh-3:konpar | -0.00814 | -0.03561 | 1.93E-02 | 1.0000 |
| 6:prukoh-3:konpar | -0.05511 | -0.09177 | -1.85E-02 | **0.0000** |
| 2:konpar-1:konpar | -0.01112 | -0.02735 | 5.12E-03 | 0.6496 |
| 4:konpar-1:konpar | -0.01171 | -0.02877 | 5.35E-03 | 0.6448 |
| 5:konpar-1:konpar | -0.01851 | -0.03689 | -1.26E-04 | **0.0461** |
| 7:konpar-1:konpar | -0.04206 | -0.06094 | -2.32E-02 | **0.0000** |
| 6:konpar-1:konpar | -0.02598 | -0.04602 | -5.94E-03 | **0.0007** |
| 3:prukoh-1:konpar | -0.02952 | -0.05036 | -8.68E-03 | **0.0001** |
| 1:prukoh-1:konpar | 0.008805 | -0.01303 | 3.06E-02 | 0.9978 |
| 2:prukoh-1:konpar | -0.02781 | -0.04806 | -7.55E-03 | **0.0002** |
| 4:prukoh-1:konpar | -0.03984 | -0.06082 | -1.89E-02 | **0.0000** |
| 5:prukoh-1:konpar | -0.01018 | -0.03574 | 1.54E-02 | 0.9982 |
| 7:prukoh-1:konpar | -0.03304 | -0.06076 | -5.31E-03 | **0.0039** |
| 6:prukoh-1:konpar | -0.08 | -0.11685 | -4.32E-02 | **0.0000** |
| 4:konpar-2:konpar | -0.00059 | -0.01753 | 1.63E-02 | 1.0000 |
| 5:konpar-2:konpar | -0.00739 | -0.02566 | 1.09E-02 | 0.9977 |
| 7:konpar-2:konpar | -0.03094 | -0.04972 | -1.22E-02 | **0.0000** |
| 6:konpar-2:konpar | -0.01486 | -0.0348 | 5.08E-03 | 0.4790 |
| 3:prukoh-2:konpar | -0.0184 | -0.03915 | 2.34E-03 | 0.1635 |
| 1:prukoh-2:konpar | 0.019921 | -0.00182 | 4.17E-02 | 0.1233 |
| 2:prukoh-2:konpar | -0.01669 | -0.03684 | 3.46E-03 | 0.2713 |
| 4:prukoh-2:konpar | -0.02873 | -0.0496 | -7.85E-03 | **0.0002** |
| 5:prukoh-2:konpar | 0.000933 | -0.02454 | 2.64E-02 | 1.0000 |
| 7:prukoh-2:konpar | -0.02192 | -0.04957 | 5.73E-03 | 0.3532 |
| 6:prukoh-2:konpar | -0.06889 | -0.10568 | -3.21E-02 | **0.0000** |
| 5:konpar-4:konpar | -0.0068 | -0.0258 | 1.22E-02 | 0.9996 |
| 7:konpar-4:konpar | -0.03035 | -0.04984 | -1.09E-02 | **0.0000** |
| 6:konpar-4:konpar | -0.01427 | -0.03488 | 6.35E-03 | 0.6288 |
| 3:prukoh-4:konpar | -0.01781 | -0.03921 | 3.59E-03 | 0.2622 |
| 1:prukoh-4:konpar | 0.020514 | -0.00184 | 4.29E-02 | 0.1220 |
| 2:prukoh-4:konpar | -0.0161 | -0.03692 | 4.73E-03 | 0.4037 |
| 4:prukoh-4:konpar | -0.02814 | -0.04966 | -6.61E-03 | **0.0006** |
| 5:prukoh-4:konpar | 0.001526 | -0.02448 | 2.75E-02 | 1.0000 |
| 7:prukoh-4:konpar | -0.02133 | -0.04947 | 6.81E-03 | 0.4444 |
| 6:prukoh-4:konpar | -0.06829 | -0.10546 | -3.11E-02 | **0.0000** |
| 7:konpar-5:konpar | -0.02355 | -0.04421 | -2.90E-03 | **0.0081** |
| 6:konpar-5:konpar | -0.00747 | -0.02919 | 1.43E-02 | 0.9998 |
| 3:prukoh-5:konpar | -0.01101 | -0.03348 | 1.15E-02 | 0.9769 |
| 1:prukoh-5:konpar | 0.027312 | 0.00393 | 5.07E-02 | **0.0055** |
| 2:prukoh-5:konpar | -0.0093 | -0.03122 | 1.26E-02 | 0.9958 |
| 4:prukoh-5:konpar | -0.02134 | -0.04392 | 1.24E-03 | **0.0922** |
| 5:prukoh-5:konpar | 0.008324 | -0.01857 | 3.52E-02 | 1.0000 |
| 7:prukoh-5:konpar | -0.01453 | -0.04349 | 1.44E-02 | 0.9705 |
| 6:prukoh-5:konpar | -0.0615 | -0.09928 | -2.37E-02 | **0.0000** |
| 6:konpar-7:konpar | 0.016084 | -0.00606 | 3.82E-02 | 0.5321 |
| 3:prukoh-7:konpar | 0.01254 | -0.01033 | 3.54E-02 | 0.9299 |
| 1:prukoh-7:konpar | 0.050865 | 0.02709 | 7.46E-02 | **0.0000** |
| 2:prukoh-7:konpar | 0.014254 | -0.00808 | 3.66E-02 | 0.7695 |
| 4:prukoh-7:konpar | 0.002215 | -0.02077 | 2.52E-02 | 1.0000 |
| 5:prukoh-7:konpar | 0.031877 | 0.004642 | 5.91E-02 | **0.0053** |
| 7:prukoh-7:konpar | 0.009024 | -0.02025 | 3.83E-02 | 1.0000 |
| 6:prukoh-7:konpar | -0.03794 | -0.07598 | 8.89E-05 | 0.0514 |
| 3:prukoh-6:konpar | -0.00354 | -0.02738 | 2.03E-02 | 1.0000 |
| 1:prukoh-6:konpar | 0.034781 | 0.010074 | 5.95E-02 | **0.0001** |
| 2:prukoh-6:konpar | -0.00183 | -0.02516 | 2.15E-02 | 1.0000 |
| 4:prukoh-6:konpar | -0.01387 | -0.03782 | 1.01E-02 | 0.8876 |
| 5:prukoh-6:konpar | 0.015793 | -0.01226 | 4.38E-02 | 0.9112 |
| 7:prukoh-6:konpar | -0.00706 | -0.0371 | 2.30E-02 | 1.0000 |
| 6:prukoh-6:konpar | -0.05403 | -0.09265 | -1.54E-02 | **0.0001** |
| 1:prukoh-3:prukoh | 0.038325 | 0.012961 | 6.37E-02 | **0.0000** |
| 2:prukoh-3:prukoh | 0.001714 | -0.02231 | 2.57E-02 | 1.0000 |
| 4:prukoh-3:prukoh | -0.01032 | -0.03495 | 1.43E-02 | 0.9964 |
| 5:prukoh-3:prukoh | 0.019338 | -0.0093 | 4.80E-02 | 0.6747 |
| 7:prukoh-3:prukoh | -0.00352 | -0.0341 | 2.71E-02 | 1.0000 |
| 6:prukoh-3:prukoh | -0.05048 | -0.08953 | -1.14E-02 | **0.0008** |
| 2:prukoh-1:prukoh | -0.03661 | -0.06149 | -1.17E-02 | **0.0000** |
| 4:prukoh-1:prukoh | -0.04865 | -0.07412 | -2.32E-02 | **0.0000** |
| 5:prukoh-1:prukoh | -0.01899 | -0.04835 | 1.04E-02 | 0.7487 |
| 7:prukoh-1:prukoh | -0.04184 | -0.0731 | -1.06E-02 | **0.0004** |
| 6:prukoh-1:prukoh | -0.08881 | -0.12839 | -4.92E-02 | **0.0000** |
| 4:prukoh-2:prukoh | -0.01204 | -0.03617 | 1.21E-02 | 0.9722 |
| 5:prukoh-2:prukoh | 0.017623 | -0.01058 | 4.58E-02 | 0.8001 |
| 7:prukoh-2:prukoh | -0.00523 | -0.03541 | 2.50E-02 | 1.0000 |
| 6:prukoh-2:prukoh | -0.0522 | -0.09093 | -1.35E-02 | **0.0003** |
| 5:prukoh-4:prukoh | 0.029662 | 0.000935 | 5.84E-02 | **0.0339** |
| 7:prukoh-4:prukoh | 0.006809 | -0.02386 | 3.75E-02 | 1.0000 |
| 6:prukoh-4:prukoh | -0.04016 | -0.07927 | -1.04E-03 | **0.0364** |
| 7:prukoh-5:prukoh | -0.02285 | -0.05682 | 1.11E-02 | 0.6817 |
| 6:prukoh-5:prukoh | -0.06982 | -0.11157 | -2.81E-02 | **0.0000** |
| 6:prukoh-7:prukoh | -0.04697 | -0.09008 | -3.86E-03 | **0.0164** |

**Table S3.** Summary statistics for anchored assembly. For each cross and for the combined pseudomolecule assembly the number of scaffolds with at least 2 markers, with at least two markers that are > 0.1 cM apart, the combined size of the anchored scaffolds, the N50, and the average coverage are shown per LG.

| LG | # scaffolds | # scaffolds >= 2 markers | # scaffolds >=2 well-spaced markers | Size (bp) | N50 (bp) | coverage |
| --- | --- | --- | --- | --- | --- | --- |
| **ParKoh** |  |  |  |  |  |  |
| 1 | 117 | 21 | 14 | 106312036 | 1301586 | 52.89792 |
| 2 | 89 | 8 | 4 | 59124686 | 886001 | 54.757 |
| 3 | 109 | 14 | 12 | 98715872 | 1184645 | 51.15361 |
| 4 | 49 | 1 | 1 | 39589978 | 1093907 | 54.17022 |
| 5 | 76 | 18 | 16 | 62735740 | 1197186 | 66.5671 |
| 6 | 47 | 7 | 6 | 19057194 | 730017 | 57.78891 |
| 7 | 45 | 6 | 5 | 38543039 | 1485176 | 60.40972 |
| X | 76 | 4 | 2 | 84017840 | 1519936 | 29.56227 |
| Sum/median | 608 | 79 | 60 | 508096385 | 1190916 | 53.41334375 |
| **KonPar** |  |  |  |  |  |  |
| 1 | 128 | 17 | 13 | 103734776 | 1143465 | 37.22859 |
| 2 | 132 | 17 | 10 | 95158272 | 1019600 | 40.18552 |
| 3 | 143 | 22 | 19 | 134318749 | 1355019 | 34.98008 |
| 4 | 109 | 24 | 12 | 110192983 | 1660236 | 40.38131 |
| 5 | 84 | 13 | 9 | 67417874 | 1136159 | 44.86084 |
| 6 | 64 | 7 | 6 | 23922075 | 733389 | 46.43391 |
| 7 | 77 | 17 | 13 | 58701654 | 1180700 | 38.07606 |
| X | 86 | 14 | 3 | 98541466 | 1540389 | 25.64274 |
| Sum/median | 823 | 131 | 85 | 691987849 | 1162083 | 38.47363125 |
| **PruKoh** |  |  |  |  |  |  |
| 1 | 50 | 5 | 3 | 46785714 | 1325001 | 41.55226 |
| 2 | 62 | 4 | 3 | 45819772 | 1106261 | 47.8152 |
| 3 | 57 | 2 | 2 | 54629290 | 1375220 | 40.13577 |
| 4 | 56 | 11 | 5 | 60827800 | 2025849 | 45.12183 |
| 5 | 33 | 3 | 2 | 20743600 | 725518 | 51.94363 |
| 6 | 14 | 0 | 0 | 7779385 | 859092 | 45.60227 |
| 7 | 27 | 4 | 2 | 17783624 | 1224531 | 49.36982 |
| X | 84 | 9 | 0 | 82253003 | 1268875 | 28.72471 |
| Sum/median | 383 | 38 | 17 | 336622188 | 1246703 | 43.78318625 |
| **Combined** |  |  |  |  |  |  |
| 1 | 167 | 43 | 30 | 117180395 | 1089296 |  |
| 2 | 170 | 29 | 17 | 101876544 | 816734 |  |
| 3 | 175 | 38 | 33 | 137279277 | 1044733 |  |
| 4 | 118 | 36 | 18 | 89703238 | 957048 |  |
| 5 | 102 | 34 | 27 | 62192538 | 880080 |  |
| 6 | 80 | 14 | 12 | 25387579 | 569059 |  |
| 7 | 88 | 27 | 20 | 52742540 | 916807 |  |
| X | 154 | 27 | 5 | 133989488 | 1249941 |  |
| Sum/median | 1054 | 248 | 162 | 720351599 | 936928 |  |

**Table S4.** Integrated AFLP and SNP map for the *L. kohalensis* x *L. paranigra* cross. The highlighted AFLP markers are located under a QTL peak in the Shaw & Lesnick 2009 study. The highlighted AFLP markers on linkage group 1 are markers where a male song and female preference QTL peak co-localize.

| scaffold | locus | LG | position (cM) | AFLP | scaffold midpoint position (bp) |
| --- | --- | --- | --- | --- | --- |
| S002761 | S002761_729410 | 1 | 0 | NA | 106423286 |
|  | as030 | 1 | 1.821 | PaggcA53 | NA |
|  | as074 | 1 | 3.282 | PggacA54 | NA |
|  | ac007 | 1 | 6.09 | PagacA52B55 | NA |
|  | ac013 | 1 | 7.301 | PcgacA51B51 | NA |
|  | ac017 | 1 | 8.604 | PgcacA07B54 | NA |
| S000817 | S000817_120415 | 1 | 9.734 | NA | 114208815 |
| S002077 | S002077_311803 | 1 | 11.288 | NA | 109788919 |
| S007909 | S007909_155126 | 1 | 13.752 | NA | 104812030 |
|  | as087 | 1 | 23.642 | PgtgcA54 | NA |
|  | as081 | 1 | 25.658 | PgtacA56 | NA |
|  | as085_x | 1 | 30.736 | PgtgcA3 | NA |
|  | as080 | 1 | 36.928 | PgtacA55 | NA |
|  | as023 | 1 | 38.868 | PaaacA63 | NA |
| S001330 | S001330_135948 | 1 | 42.507 | NA | 100949675 |
| S001771 | S001771_116507 | 1 | 46.906 | NA | 105601414 |
| S000392 | S000392_74030 | 1 | 51.737 | NA | NA |
| S001680 | S001680_315523 | 1 | 52.615 | NA | 88172650 |
| S000409 | S000409_474112 | 1 | 53.517 | NA | 84174036 |
| S001489 | S001489_769426 | 1 | 56.166 | NA | 41585373 |
| S004205 | S004205_29098 | 1 | 57.827 | NA | 50498860 |
| S000949 | S000949 205067 | 1 | 58.259 | NA | 42520204 |
| S008139 | S008139 68543 | 1 | 58.542 | NA | NA |
| S000696 | S000696 137337 | 1 | 58.549 | NA | NA |
| S002946 | S002946 738803 | 1 | 58.707 | NA | 67349798 |
| C120306 | C120306 385 | 1 | 58.847 | NA | NA |
| S006572 | S006572 95801 | 1 | 58.928 | NA | 46181571 |
| S001914 | S001914 404347 | 1 | 59.118 | NA | 66160290 |
| S009296 | S009296 110864 | 1 | 59.329 | NA | 65039174 |
| S000663 | S000663 611964 | 1 | 59.641 | NA | 53947499 |
| S004747 | S004747 292840 | 1 | 59.784 | NA | 49933664 |
| S000671 | S000671 1033505 | 1 | 61.079 | NA | NA |
| S001489 | S001489 639186 | 1 | 61.694 | NA | 41585373 |
| S004313 | S004313 69835 | 1 | 61.699 | NA | 40812282 |
| S002548 | S002548 423714 | 1 | 62.493 | NA | 45749393 |
| S000105 | S000105 348801 | 1 | 63.591 | NA | 36945493 |
| S004771 | S004771 628132 | 1 | 64.917 | NA | 31081394 |
| S004771 | S004771 1175996 | 1 | 65.21 | NA | 31081394 |
| S002151 | S002151 1377807 | 1 | 66.292 | NA | 22870317 |
|  | as034 | 1 | 68.191 | PatgcA52 | NA |
|  | as012 | 1 | 70.107 | PccacA55 | NA |
|  | ac014 | 2 | 0 | PgaacA10B60 | NA |
| S001206 | S001206 1546586 | 2 | 2.303 | NA | 1821533 |
| S000518 | S000518 766492 | 2 | 6.245 | NA | 6192438 |
|  | as077 | 2 | 26.628 | PgggcA52 | NA |
| S003191 | S003191 528616 | 2 | 31.612 | NA | 16874625 |
| S001838 | S001838 6021 | 2 | 42.989 | NA | 23233385 |
| S000416 | S000416 552586 | 2 | 47.133 | NA | 24694735 |
| S004218 | S004218 23553 | 2 | 51.627 | NA | 31927917 |
| S001550 | S001550 214202 | 2 | 52.41 | NA | 33996652 |
| S003798 | S003798 463488 | 2 | 53.5 | NA | 36725738 |
| S002376 | S002376 431585 | 2 | 54.951 | NA | 38303813 |
|  | as052 | 2 | 57.787 | PcggcA53 | NA |
| S005289 | S005289 526503 | 2 | 58.474 | NA | 45049140 |
| S000230 | S000230 200879 | 2 | 58.756 | NA | 46073354 |
| S002156 | S002156 413885 | 2 | 61.643 | NA | 90147661 |
| S003079 | S003079 192141 | 2 | 61.705 | NA | 56392065 |
| S004728 | S004728 52289 | 2 | 61.99 | NA | 77439224 |
| S001050 | S001050 43796 | 2 | 62.025 | NA | 71210412 |
| S001881 | S001881 642832 | 2 | 62.193 | NA | 60129008 |
| S003118 | S003118 235274 | 2 | 62.379 | NA | 56724422 |
| S003735 | S003735 82587 | 2 | 62.896 | NA | NA |
|  | ac006 | 2 | 63.793 | PacacA56B69 | NA |
|  | as040 x | 2 | 64.823 | PcagcA08 | NA |
| S003067 | S003067_20757 | 2 | 67.28 | NA | 80842581 |
| S000199 | S000199_193260 | 2 | 74.457 | NA | 83353553 |
| S001797 | S001797_1827615 | 2 | 75.671 | NA | 85052994 |
| S001855 | S001855_342231 | 2 | 82.018 | NA | 89412353 |
| S004792 | S004792_152639 | 2 | 85.245 | NA | NA |
| S001901 | S001901_315332 | 2 | 87.985 | NA | 94264609 |
| S001602 | S001602_266912 | 2 | 109.791 | NA | 99970919 |
| S000793 | S000793_627068 | 2 | 116.258 | NA | NA |
|  | as115 | 3 | 0 | PttacA54 | NA |
|  | as083 | 3 | 3.848 | PgtacA58 | NA |
| S001338 | S001338_29965 | 3 | 5.698 | NA | 168668 |
|  | as037 | 3 | 9.753 | PcaacA57 | NA |
| S000075 | S000075_156551 | 3 | 11.248 | NA | 1642392 |
| S002528 | S002528_794391 | 3 | 19.911 | NA | 18001031 |
| S009989 | S009989_70944 | 3 | 21.779 | NA | 6045815 |
| S002528 | S002528_794393 | 3 | 23.255 | NA | 18001031 |
| S005403 | S005403_7134 | 3 | 27.914 | NA | 18662141 |
|  | ac009 a | 3 | 48.316 | PaggcA07B19 | NA |
|  | as063 | 3 | 49.481 | PgaacA60 | NA |
|  | as064 | 3 | 51.877 | PgaacA61 | NA |
|  | as025 | 3 | 53.754 | PaagcA57 | NA |
| S000558 | S000558_1959639 | 3 | 59.995 | NA | 40534699 |
| S000558 | S000558_1232157 | 3 | 60.337 | NA | 40534699 |
|  | as088 | 3 | 63.867 | PgtgcA55 | NA |
|  | as069 | 3 | 64.465 | PgcacA51 | NA |
|  | ad100.as079 | 3 | 65.149 | NA | NA |
| S004777 | S004777_221108 | 3 | 66.442 | NA | 36430961 |
| S000558 | S000558_748836 | 3 | 66.865 | NA | 40534699 |
| S007419 | S007419_788393 | 3 | 67.162 | NA | 42061284 |
| S002934 | S002934_242739 | 3 | 68.016 | NA | 56760880 |
| S003072 | S003072_393178 | 3 | 68.244 | NA | 63064570 |
| S001785 | S001785_133632 | 3 | 68.301 | NA | 60906412 |
| S000726 | S000726_1782089 | 3 | 68.399 | NA | 48678466 |
| S000529 | S000529_530363 | 3 | 68.572 | NA | 72186930 |
|  | as108 a | 3 | 68.59 | PtgacA03 | NA |
| S002665 | S002665_282741 | 3 | 68.951 | NA | NA |
| S005483 | S005483_46807 | 3 | 69.079 | NA | 62650842 |
| S013086 | S013086_40139 | 3 | 69.12 | NA | NA |
| S002194 | S002194_96231 | 3 | 69.157 | NA | 66669112 |
| S006750 | S006750_274965 | 3 | 69.245 | NA | NA |
| S004654 | S004654_237626 | 3 | 69.303 | NA | NA |
| S002002 | S002002_376411 | 3 | 69.412 | NA | 44975632 |
| S003472 | S003472_9538 | 3 | 70.555 | NA | 82404642 |
| S002613 | S002613_31096 | 3 | 71.526 | NA | NA |
| S001060 | S001060_728231 | 3 | 72.259 | NA | 89191537 |
| S002297 | S002297_775779 | 3 | 72.322 | NA | 91133138 |
| S001060 | S001060_1929755 | 3 | 72.461 | NA | 89191537 |
| S006865 | S006865_338439 | 3 | 72.867 | NA | 101275163 |
|  | as101 | 3 | 73.62 | PtcgcA09 | NA |
| S001265 | S001265_211884 | 3 | 74.555 | NA | 106649134 |
| S000385 | S000385_1480834 | 3 | 75.848 | NA | 108764126 |
| S001106 | S001106_481878 | 3 | 82.071 | NA | 114989880 |
|  | as114_ax | 3 | 86.713 | PttacA02 | NA |
| S001275 | S001275_1127865 | 3 | 88.98 | NA | 122746700 |
| S002311 | S002311_254807 | 3 | 89.321 | NA | 123571440 |
|  | as076_a | 4 | 0 | PgggcA01 | NA |
|  | as073_a | 4 | 1.753 | PggacA01 | NA |
| S005844 | S005844_224400 | 4 | 18.591 | NA | 4097279 |
|  | as117_a | 4 | 41.053 | PttacA06 | NA |
|  | as099 | 4 | 41.551 | PtcacA54 | NA |
| S001783 | S001783_279356 | 4 | 43.04 | NA | 63112104 |
| S000590 | S000590_1997032 | 4 | 43.249 | NA | 29331168 |
| S014891 | S014891_36875 | 4 | 43.589 | NA | NA |
| S003206 | S003206_599818 | 4 | 43.711 | NA | NA |
| S000836 | S000836_228275 | 4 | 43.801 | NA | NA |
| S000679 | S000679_623688 | 4 | 43.895 | NA | 57825201 |
|  | as075_a | 4 | 44.445 | PggacA13 | NA |
| S002196 | S002196_584574 | 4 | 44.674 | NA | 42514727 |
| S001005 | S001005_1065597 | 4 | 44.857 | NA | 39878575 |
| S003635 | S003635_7993 | 4 | 45.443 | NA | 63978342 |
| S009873 | S009873_154713 | 4 | 49.074 | NA | 71866807 |
| S002058 | S002058_706797 | 4 | 49.408 | NA | 72523035 |
| S000455 | S000455_692625 | 4 | 52.283 | NA | 73544308 |
|  | as029_a | 4 | 61.5 | PagacA5 | NA |
| S001486 | S001486_61579 | 4 | 64.17 | NA | 77623428 |
|  | as028_x | 4 | 65.829 | PagacA1 | NA |
| S001279 | S001279_277406 | 4 | 67.269 | NA | 78135382 |
| S001608 | S001608_399358 | 4 | 83.462 | NA | 89198891 |
|  | as021 | 5 | 0 | PaaacA54 | NA |
| S005326 | S005326_57790 | 5 | 1.807 | NA | 466530 |
|  | ac016 | 5 | 2.394 | PgagcA56B62 | NA |
|  | ac011 | 5 | 4.24 | PcagcA52B53 | NA |
|  | ac015 | 5 | 5.865 | PgaacA62B66 | NA |
|  | as116_a | 5 | 7.016 | PttacA05 | NA |
| S002190 | S002190_273188 | 5 | 13.671 | NA | 3452268 |
|  | as100 | 5 | 15.717 | PtcgcA54 | NA |
| S000809 | S000809_365747 | 5 | 19.221 | NA | 6005121 |
|  | as066_a | 5 | 21.423 | PgagcA08 | NA |
|  | as027_a | 5 | 25.245 | PacacA06 | NA |
|  | ad082.as062 | 5 | 26.337 | NA | NA |
|  | as045 | 5 | 27.055 | PccacA53 | NA |
| S004462 | S004462_54127 | 5 | 28.207 | NA | 13078451 |
| S005610 | S005610_84211 | 5 | 28.338 | NA | 13179099 |
| S000366 | S000366_42046 | 5 | 28.803 | NA | 17964325 |
| S000745 | S000745_339612 | 5 | 30.425 | NA | 21039003 |
| S002565 | S002565_116928 | 5 | 33.166 | NA | 22902902 |
| S004683 | S004683_48328 | 5 | 34.768 | NA | 24389284 |
| S006506 | S006506_7146 | 5 | 35.301 | NA | 27839633 |
| S005519 | S005519_111372 | 5 | 37.975 | NA | 32385211 |
| S000305 | S000305_70577 | 5 | 40.26 | NA | 33073185 |
| S000979 | S000979_164421 | 5 | 41.485 | NA | 35633669 |
| S005459 | S005459_70152 | 5 | 42.698 | NA | 37847956 |
| S000180 | S000180_656202 | 5 | 44.503 | NA | 40272446 |
| S021890 | S021890_296 | 5 | 45.368 | NA | 42534517 |
|  | as068 | 5 | 46.655 | PgagcA51 | NA |
| S005334 | S005334_732385 | 5 | 48.033 | NA | 43080568 |
| S005064 | S005064_175253 | 5 | 49.954 | NA | 45035858 |
|  | as091 | 5 | 52.052 | PtaacA51 | NA |
| S003838 | S003838_738497 | 5 | 54.849 | NA | 46738481 |
| S001560 | S001560_398299 | 5 | 59.376 | NA | 50508185 |
| S004681 | S004681_23198 | 5 | 61.431 | NA | 53592894 |
|  | as057 | 5 | 62.937 | PctacA55 | NA |
|  | as058 | 5 | 63.812 | PctacA56 | NA |
|  | as059 | 5 | 65.211 | PctacA57 | NA |
|  | as043 | 5 | 71.183 | PcagcA51 | NA |
|  | as118_x | 5 | 72.676 | PttacA13 | NA |
|  | as041 | 5 | 73.969 | PcagcA56 | NA |
| S007270 | S007270_308120 | 5 | 76.332 | NA | 58235091 |
| S002503 | S002503_741242 | 5 | 77.495 | NA | 58956874 |
|  | as110 | 6 | 0 | PtggcA52 | NA |
|  | as047_x | 6 | 1.51 | PcgacA02 | NA |
| S001034 | S001034_78522 | 6 | 2.72 | NA | 24400658 |
| S022584 | S022584_1139 | 6 | 5.895 | NA | 23964288 |
|  | as105 | 6 | 6.664 | PtgacA58 | NA |
| S005236 | S005236_14409 | 6 | 7.404 | NA | 16442306 |
| S001507 | S001507_262717 | 6 | 8.078 | NA | 13978219 |
| S002799 | S002799_37080 | 6 | 10.159 | NA | 22063251 |
|  | ac003 | 6 | 11.251 | PaagcA61B63 | NA |
|  | ac019 | 6 | 13.745 | PtgacA57B60 | NA |
| S001904 | S001904_134885 | 6 | 15.337 | NA | 16620760 |
|  | as109 | 6 | 17.735 | PtgacA56 | NA |
|  | as039 | 6 | 20.554 | PcagcA54 | NA |
| S000218 | S000218_4280 | 6 | 21.865 | NA | 10002544 |
| S001761 | S001761_1745572 | 6 | 27.303 | NA | 6635712 |
| S005439 | S005439_103276 | 6 | 35.966 | NA | 4315651 |
| S011721 | S011721_76537 | 6 | 47.364 | NA | 849742 |
| S003103 | S003103_949190 | 7 | 0 | NA | 1904957 |
| S008107 | S008107_29859 | 7 | 3.613 | NA | 2842152 |
| S003103 | S003103_1019194 | 7 | 3.994 | NA | 1904957 |
| S000557 | S000557_589404 | 7 | 7.085 | NA | 3829489 |
| S002304 | S002304_728504 | 7 | 9.873 | NA | 4666925 |
| S002736 | S002736_346176 | 7 | 10.997 | NA | 5494164 |
| S002736 | S002736_115345 | 7 | 12.279 | NA | 5494164 |
| S014172 | S014172_53634 | 7 | 13.346 | NA | 5971292 |
| S000628 | S000628_1159720 | 7 | 17.535 | NA | 8386281 |
| S000628 | S000628_169050 | 7 | 18.732 | NA | 8386281 |
|  | as033 | 7 | 21.278 | PatacA55 | NA |
|  | as071_a | 7 | 22.74 | PgcgcA11 | NA |
| S000677 | S000677_527760 | 7 | 24.152 | NA | 12625625 |
|  | as070_a | 7 | 26.358 | PgcacA10 | NA |
| S002087 | S002087_530732 | 7 | 44.525 | NA | 31673319 |
| S004909 | S004909_906373 | 7 | 50.728 | NA | 39335421 |
| S010292 | S010292_145463 | 7 | 53.418 | NA | 40115847 |
| S004220 | S004220_129736 | 7 | 53.983 | NA | 43031005 |
| S001469 | S001469_584893 | 7 | 65.118 | NA | 50836753 |
| S002493 | S002493_175861 | 7 | 67.129 | NA | 48491131 |
|  | xs032_x | X | 0 | PttacA04 | NA |
| S000571 | S000571_30159 | X | 2.403 | NA | 4915702 |
|  | xd024 | X | 11.115 | PtagcB55 | NA |
| S001780 | S001780_509180 | X | 24.446 | NA | 10902308 |
|  | xd004 | X | 25.132 | PatacB52 | NA |
|  | xs020 | X | 29.168 | PatgcA56 | NA |
| S000766 | S000766 827724 | X | 30.52 | NA | 11869181 |
| S000360 | S000360 70566 | X | 32.459 | NA | 17072014 |
| S003455 | S003455 1122569 | X | 35.526 | NA | 19186595 |
| S004887 | S004887 81621 | X | 42.013 | NA | NA |
|  | xs024 x | X | 43.011 | PcggcA10 | NA |
| S000604 | S000604 43741 | X | 43.459 | NA | NA |
| S000648 | S000648 2315784 | X | 44.883 | NA | 34819414 |
| S000219 | S000219 2024096 | X | 45.729 | NA | 26925418 |
| S000777 | S000777 1339236 | X | 46.733 | NA | 30344332 |
| S000777 | S000777 461461 | X | 46.887 | NA | 30344332 |
| S000648 | S000648 2255296 | X | 47.453 | NA | 34819414 |
| S001873 | S001873 641123 | X | 49.384 | NA | 36757980 |
|  | xd016 x | X | 49.672 | PatgcB02 | NA |
| S001241 | S001241 171252 | X | 53.266 | NA | 44632865 |
| S003053 | S003053 264327 | X | 55.937 | NA | 56937852 |
| S000327 | S000327 278424 | X | 56.293 | NA | 55828014 |
| S000470 | S000470 76715 | X | 56.911 | NA | 59714238 |
|  | xd008 x | X | 57.404 | PgggcB22 | NA |
| S003307 | S003307 279141 | X | 57.747 | NA | 63486064 |
| S001912 | S001912 587919 | X | 58.069 | NA | 62108629 |
| S000965 | S000965 1228070 | X | 59.663 | NA | 73444967 |
| S000808 | S000808 1583160 | X | 59.672 | NA | 69248171 |
| S006304 | S006304 697561 | X | 59.908 | NA | NA |
| S013985 | S013985 11114 | X | 60.241 | NA | 82920002 |
|  | xd020 | X | 63.263 | PcggcB53 | NA |
| S007907 | S007907 144642 | X | 64.648 | NA | 94604867 |
|  | xs011 | X | 67.511 | PaaacA60 | NA |
| S001930 | S001930 1745657 | X | 68.641 | NA | 88122926 |
| S001930 | S001930 2410442 | X | 69.247 | NA | 88122926 |
| S004832 | S004832 609791 | X | 70.344 | NA | 91521965 |
| S001247 | S001247 68215 | X | 71.108 | NA | 92776928 |
|  | xs012 | X | 75.227 | PaagcA53 | NA |
| S000238 | S000238 1169086 | X | 90.242 | NA | 113789231 |
| S003071 | S003071 613045 | X | 95.05 | NA | 116953618 |
| S002737 | S002737 694631 | X | 117.43 | NA | NA |
| S000176 | S000176 35913 | X | 126.585 | NA | 132559092 |
| S001187 | S001187 211572 | X | 130.879 | NA | 127829472 |
| S003230 | S003230 1425064 | X | 131.437 | NA | 129558233 |
| S002008 | S002008 1089204 | X | 131.468 | NA | 126707516 |

**Figure S1.**
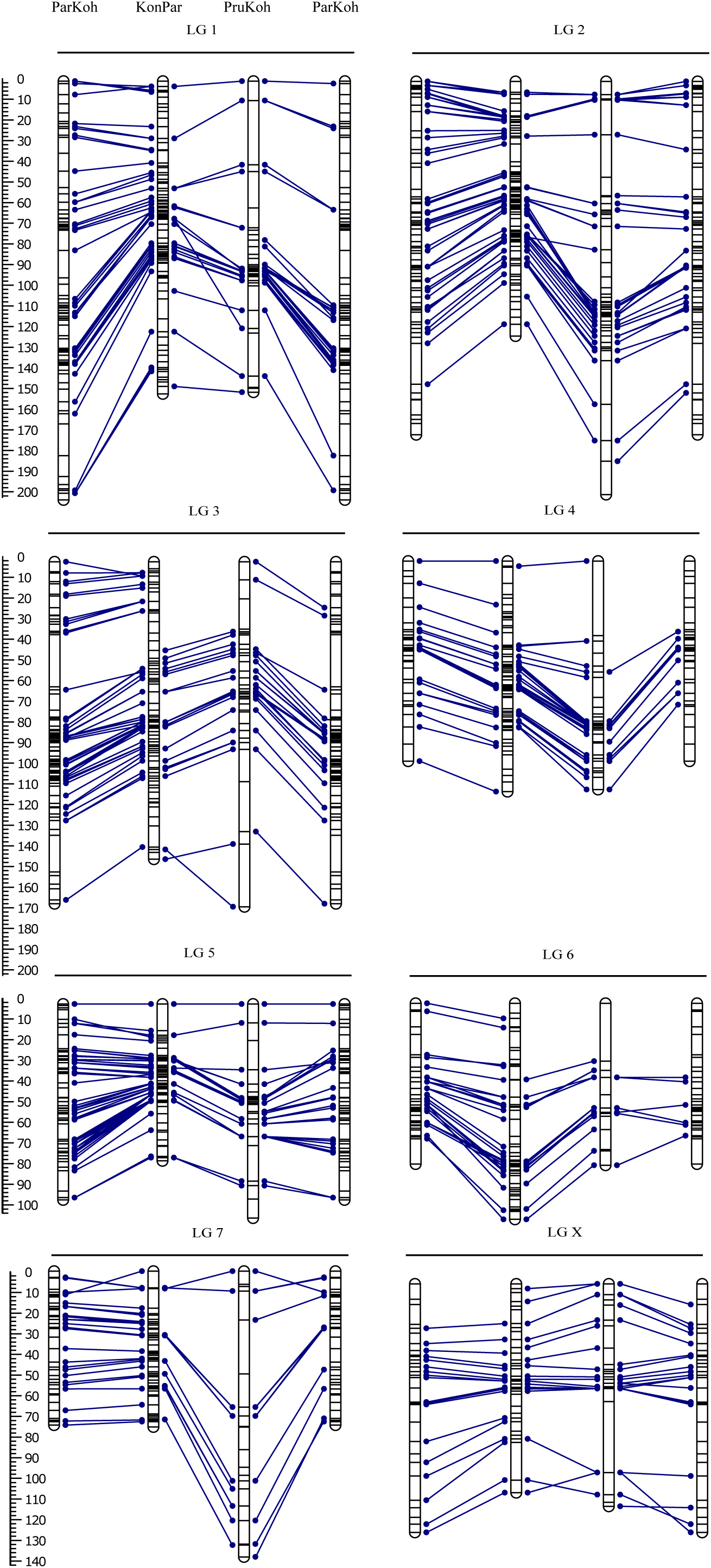
Comprehensive linkage maps. Bars represent linkage groups (LG) for ParKoh, KonPar, and PruKoh. Lines within the bars indicate marker positions. The scale on the left gives marker position in cM. Blue lines connect markers on the same scaffold between the different maps (homologous markers). The map for ParKoh is shown twice to facilitate comparisons across all three maps.

**Figure S2.**
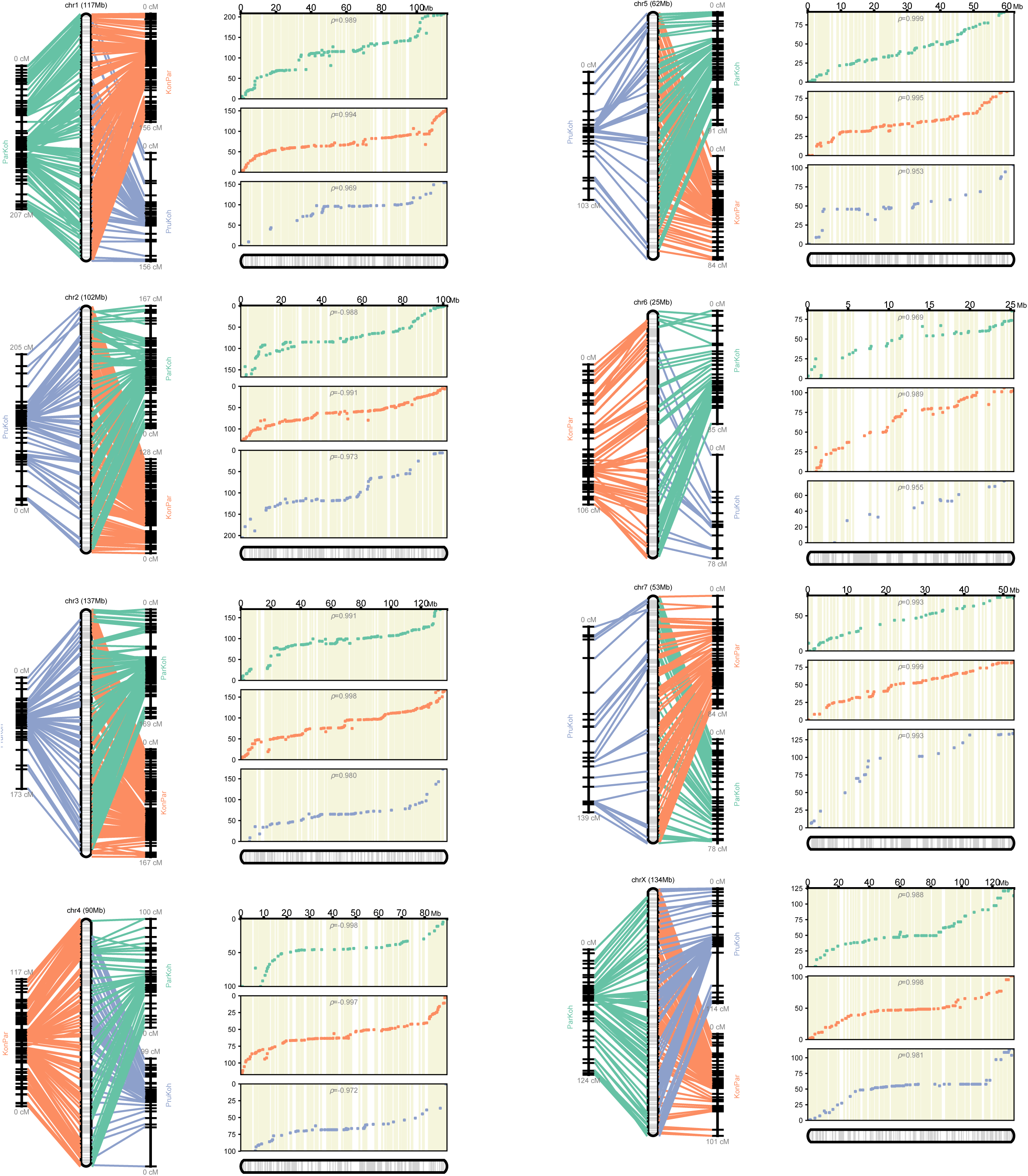
ALLMAPS output. For each of the linkage groups (chr) the relative order with respect to the shared map (i.e. the pseudomolecule assembly) is shown as well as Spearman’s rho (ρ) for the strength of correlation between marker orders.

**Figure S3.**
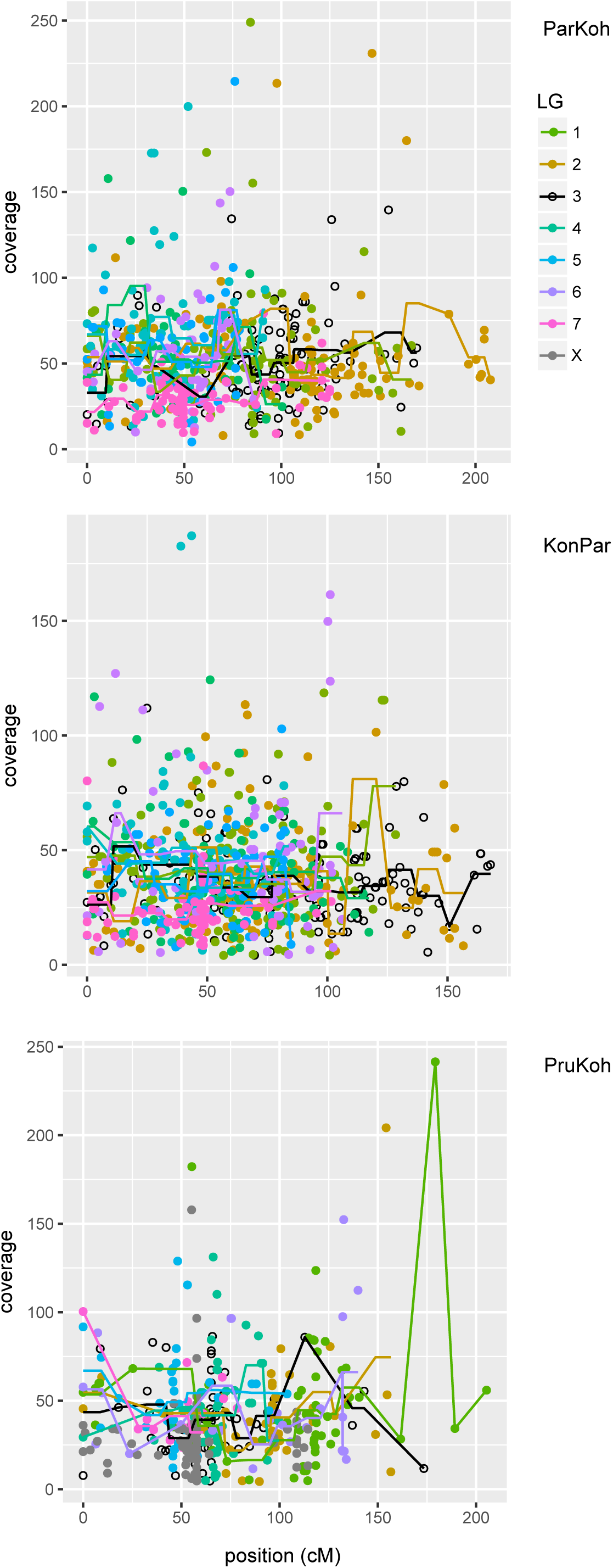
Coverage per cross per linkage group. For each of the three linkage maps (ParKoh, KonPar, PruKoh) the variation in coverage across the 8 linkage groups is shown. Coverage is calculated as the average (across individuals) read count per marker (points). Solid lines show 10-cM non-sliding window averages.

